# Modulation of bacterial multicellularity via spatiotemporal polysaccharide secretion

**DOI:** 10.1101/2020.02.20.957654

**Authors:** Salim T. Islam, Israel Vergara Alvarez, Fares Saïdi, Annick Giuseppi, Evgeny Vinogradov, Castrese Morrone, Gael Brasseur, Gaurav Sharma, Anaïs Benarouche, Jean-Luc Bridot, Gokulakrishnan Ravicoularamin, Alain Cagna, Charles Gauthier, Mitchell Singer, Henri-Pierre Fierobe, Tâm Mignot, Emilia M. F. Mauriello

## Abstract

The development of multicellularity is a key evolutionary transition allowing for differentiation of physiological functions across a cell population that confers survival benefits; among unicellular bacteria, this can lead to complex developmental behaviours and the formation of higher-order community structures. Herein, we demonstrate that in the social δ-proteobacterium *Myxococcus xanthus*, the secretion of a novel secreted biosurfactant polysaccharide (BPS) is temporally and spatially modulated within communities, mediating swarm migration as well as the formation of multicellular swarm biofilms and fruiting bodies. BPS is a type IV pilus-inhibited acidic polymer built of randomly-acetylated β-linked tetrasaccharide repeats. Both BPS and the “shared good” EPS are produced by dedicated Wzx/Wzy-dependent polysaccharide assembly pathways distinct from that responsible for spore coat assembly. To our knowledge, such pathways have never-before been explicitly shown to synthesize a biosurfactant. Together, these data reveal the central role of secreted polysaccharides in the intricate behaviours coordinating bacterial multicellularity.

## Introduction

Multicellularity is denoted by the differentiation of physiological functions across a contiguous cell population, with its development regarded as a key evolutionary transition [1]. To attain this level of organizational complexity, cells generally must be able to proliferate, specialize, communicate, interact, and move, with these behaviours promoting an increase in the size of a cell collective and the development of higher-order structures [2]. Though typically associated with metazoan organisms, multicellular physiology is also displayed by bacteria, with the best-studied examples being the formation of biofilms and fruiting bodies [3–6].

Secreted long-chain polysaccharides are an important mediator of multicellularity as they serve to retain and organize cells as well as to physically and biochemically buffer the community within the context of an extracellular matrix [7], thus enhancing survival and fitness. Monospecies bacterial biofilms have thus been intensively studied with respect to their effects on inter-cell communication, leading to differences in gene regulation and changes in matrix polysaccharide production. However, in-depth knowledge of the mechanisms used by bacteria to modulate multicellular physiology in such communities is limited.

Due to its complex social predatory lifecycle, the Gram-negative δ-proteobacterium *Myxococcus xanthus* has emerged as a leading model system in which to simultaneously study multiple factors contributing to organizational complexity. This soil bacterium is capable of saprophytic feeding on products derived from predation of other bacteria [8]. Two forms of motility are required for this complex physiology: type IV pilus (T4P)-dependent group (i.e. “social” [S]) motility [9, 10] on soft surfaces, and single-cell gliding (i.e. “adventurous” [A]) motility on hard surfaces mediated by directed transport and substratum coupling of the Agl–Glt trans-envelope complex [11, 12]. Upon local nutrient depletion, cells initiate a developmental cycle resulting in aggregation and fruiting body formation within 72 h, generating three differentiated cell subpopulations: (i) cells that form desiccation-resistant myxospores in the centre of the fruiting body, (ii) those that remain at the base of the fruiting body, termed “peripheral rods”, and (iii) forager cells that continue their outward motility away from the fruiting body [13].

*M. xanthus* produces several known long-chain polysaccharides that are central to its complex lifecycle. In addition to the O-antigen polymer that caps its LPS and is implicated in motility [14–16], *M. xanthus* biosynthesizes a poorly characterized “slime” polysaccharide that facilitates adhesion of the Glt gliding motility complex proteins to the substratum, and is deposited in trails behind surface-gliding cells [17, 18]. <u>E</u>xo<u>p</u>oly<u>s</u>accharide (EPS) is a specific secreted polymer of this bacterium that is important for T4P-dependent swarm spreading; it is also crucial for biofilm formation as it constitutes a large portion of the extracellular matrix in stationary *M. xanthus* biofilms and connects cells via a network of fibrils [19–21]. The production of EPS requires the presence of a T4P [22], which affects the Dif chemosensory pathway (reviewed elsewhere [13]). Finally, cells undergoing sporulation synthesize the <u>ma</u>jor <u>s</u>pore <u>c</u>oat (MASC) polymer that surrounds myxospores [23, 24].

The most widespread polysaccharide biosynthesis paradigm is the flippase/polymerase (Wzx/Wzy)-dependent pathway [25]. It is used by Gram-negative and Gram-positive bacteria as well as Archaea to produce a wide range of secreted and/or cell surface-associated bacterial polymers [26] including capsular polysaccharide, adhesive hold-fast polymer, spore-coat polymer, O-antigen, and exopolysaccharide [27, 28]. Wzx/Wzy-dependent polysaccharide assembly is a complex process [29] involving a suite of integral inner-membrane (IM) proteins containing multiple α-helical transmembrane segments (TMS) [30]. At the cytoplasmic leaflet of the IM, individual polysaccharide repeat units are built on an undecaprenyl pyrophosphate (UndPP) carrier. UndPP-linked repeats are then translocated across the IM by the Wzx flippase [31, 32] via a putative antiport mechanism [33, 34]. Defects in this step, resulting in a buildup of UndPP-linked repeat units for a given pathway, can have adverse effects on cell growth as well as polysaccharide synthesis by other pathways in the same cell, all dependent on UndPP-linked sugars [35]. Once in the periplasmic leaflet of the IM, repeat units are joined together by two key periplasmic loops of the Wzy polymerase [36–39], resulting in polymer extension at the reducing terminus of the growing chain [40]. Repeat-polymerization defects in a given pathway requiring UndPP may also affect other pathways requiring UndPP due to sequestration of the cellular UndPP pool. Associated polysaccharide co-polymerase (PCP) proteins determine the modal lengths for the growing polymer; these proteins typically contain two TMS with a large intervening coiled-coil periplasmic domain. Wzc proteins of the PCP-2A class (typically from Gram-negative species) further contain a cytosolic bacterial tyrosine autokinase (BY-kinase) domain fused to the 2^nd^ TMS of the PCP, whereas PCP-2B Wzc proteins (largely from Gram-positive species) are phosphorylated by a separately-encoded Wze tyrosine kinase [41]. For Gram-negative bacteria, once a polymer has been synthesized in the periplasm, it is then secreted outside the cell through the periplasm-spanning Wza translocon embedded in the OM [42, 43] (**Fig 1A**). Wzc proteins have also been implicated in polymer secretion likely through contacts formed with their cognate Wza translocon [41].

**Figure 1.**
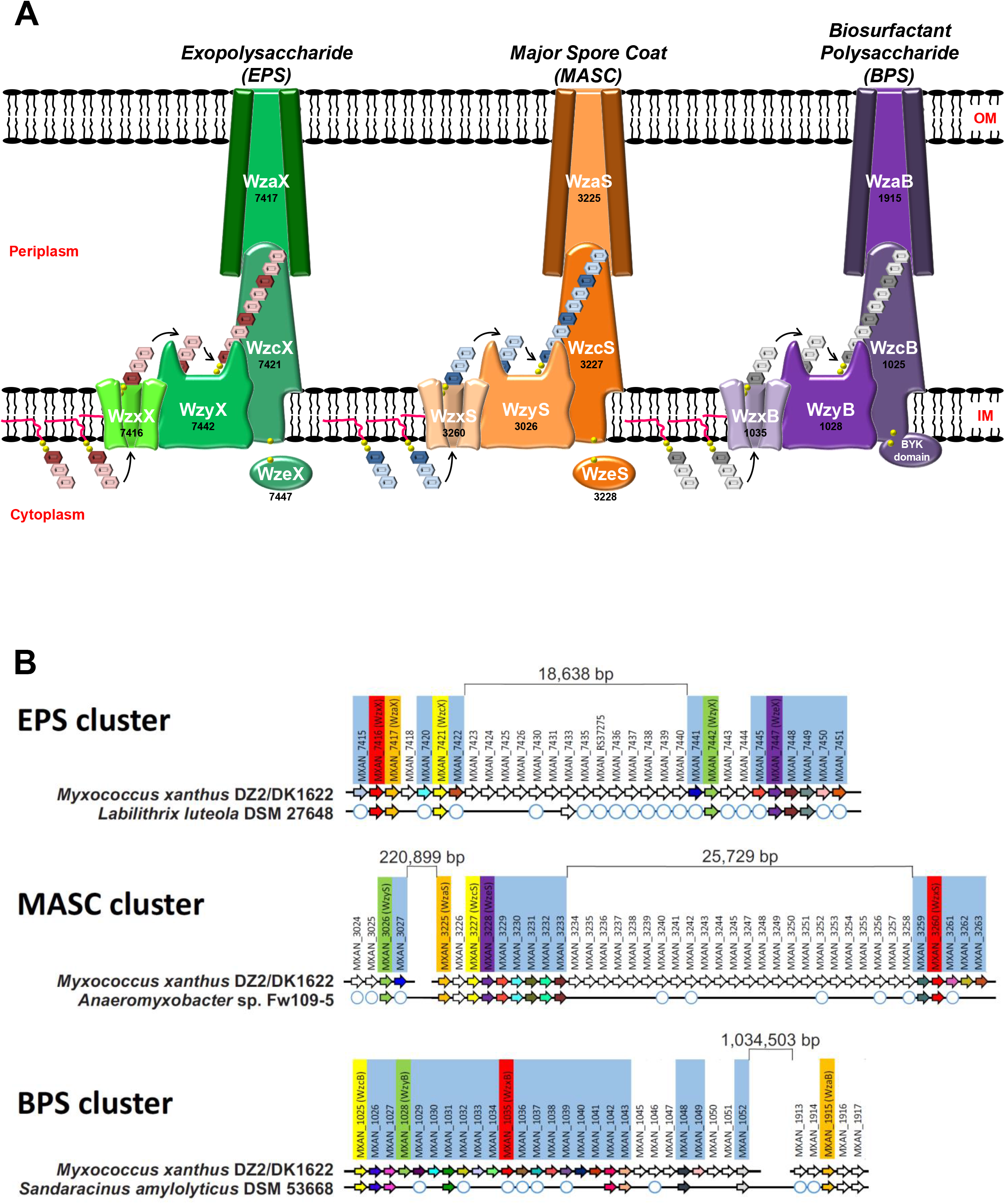
Wzx/Wzy-dependent polysaccharide biosynthesis pathways encoded by *M. xanthus*. **(A)** Schematic representation of the three Wzx/Wzy-dependent polysaccharide assembly pathways in *M. xanthus* DZ2. The genomic “MXAN” locus tag identifier for each respective gene (*black*) has been indicated below the specific protein name (*white*). **(B)** Gene conservation and synteny diagrams for EPS, MASC, and BPS clusters in *M. xanthus* DZ2 compared to the evolutionarily closest genome containing a contiguous cluster (see **S1 Fig**). Locus tags highlighted by pale blue boxes correspond to genes such as enzymes involved in monosaccharide synthesis, modification, or incorporation into precursor repeat units of the respective polymer. White circles depict the presence of a homologous gene encoded elsewhere in the chromosome (but not syntenic with the remainder of the EPS/MASC/BPS biosynthesis cluster).

The *M. xanthus* genome encodes proteins that constitute multiple, yet incompletely annotated, Wzx/Wzy-dependent polysaccharide assembly pathways. The first of these pathways is responsible for EPS biosynthesis[44], while the second synthesizes the MASC polymer that surrounds myxospores [45, 46]. Disparate Wzx/Wzy-dependent pathway proteins seemingly not involved in either EPS or MASC biosynthesis have also each been studied in isolation [17, 47], hinting at the possibility of a third such assembly pathway in *M. xanthus* for which the product, as well as the importance of its interplay with other polymers, are entirely unknown.

Herein, we describe the constituents of a newly-identified Wzx/Wzy-dependent polysaccharide assembly pathway in *M. xanthus*, as well as previously-unknown [44–46] EPS- and MASC-pathway components; this new third pathway is revealed to synthesize a T4P-regulated novel <u>b</u>iosurfactant <u>p</u>oly<u>s</u>accharide (BPS) that exerts direct effects on swarm-level *M. xanthus* behaviours. Spatiotemporal modulation of EPS and BPS pathways within *M. xanthus* communities is shown to be important for migration, development, and predation. This illustrates the importance of differentially-regulated polysaccharide production for complex, coordinated multicellular behaviours in bacteria.

## Results

### *M. xanthus* encodes three complete Wzx/Wzy-dependent polysaccharide biosynthesis pathways

To identify the core assembly-and-export constituents of each pathway, proteins with Pfam domains attributed to Wzx (domain: Polysacc_synt), Wzy (domain: PF04932 [Wzy]), Wzc (domain: PF02706 [Wzz]), and Wza (domain: Poly_export) were first identified in the *M. xanthus* genome [48]. Protein hits and other cluster constituents were further subjected to fold-recognition comparisons against existing protein structures in the PDB. These data, combined with TMS predictions and intra-protein pairwise alignments, were used to annotate the core components of each of the three assembly pathways (**Fig 1A, S2 Table**). Genes in the EPS assembly pathway and developmental MASC assembly pathway — (re)named according to longstanding convention [49, 50] and given suffixes “X” and “S” to denote e<u>x</u>opolysaccharide and <u>s</u>pore coat, respectively — were detected in clusters, in-line with previous reports [44, 46] (**S1A,B Fig**). Consistent with the high molecular-weight polysaccharide-assembly function of Wzx/Wzy-dependent pathways in Gram-negative and Gram-positive bacteria, numerous predicted glycosyltransferase (GTase) proteins were also found encoded in the immediate vicinity of the identified polymer assembly proteins (**Fig 1B, S3 Table**). In addition to the EPS and MASC assembly clusters, we further identified a novel third gene cluster encoding Wzx/Wzy-dependent pathway proteins (given suffixes “B” to denote <u>b</u>iosurfactant, see below) (**S1C Fig**). Similar to the EPS and MASC clusters, this third cluster was also highly enriched for genes encoding potential GTases (**S3 Table**), consistent with this third Wzx/Wzy-dependent pathway also producing a high molecular-weight polysaccharide (**Fig 1A**). As Wzx proteins are exquisitely-specific to the structure of individual UndPP-linked repeats [31, 35], the identification of three distinct Wzx flippases (along with their respective GTase-containing gene clusters) is indicative of the production of three structurally-distinct high molecular-weight polysaccharides by such assembly pathways in *M. xanthus*.

Though Wzx/Wzy-dependent pathway proteins are typically encoded in contiguous gene clusters [27, 51], each of the three assembly pathways in *M. xanthus* is encoded at a minimum of two separated chromosomal loci. However, homology studies of related genomes revealed syntenic and contiguous clusters in related bacteria, helping to reconcile the various insertions. The *M. xanthus* EPS cluster was found to contain an 18.739 kbp insertion separating the upstream half (encoding WzxX, WzaX, and WzcX) from the downstream half (encoding WzyX and WzeX); however, a contiguous version of this cluster was detected in the genome of *Labilithrix luteola* (**Fig 1B, S1A Fig**). Even larger insertions were found in the *M. xanthus* MASC cluster, with the tract encoding WzyS separated from the tract encoding WzaS, WzcS, and WzeS by 223.323 kbp; a 25.729 kbp insertion was also identified between the tract encoding WzxS and the WzaS-WzcS-WzeS-encoding segment. Yet, a contiguous MASC assembly cluster lacking the intervening genes was detected in *Anaeromyxobacter* sp. Fw109-5 (**Fig 1B, S1B Fig**).

Genes encoding assembly proteins WzcB, WzyB, and WzxB were located close to each other on the *M. xanthus* chromosome, interspersed with putative glycosyltransferase genes, denoting the presence of a potential BPS cluster; however, no gene encoding a WzaB protein to complete this new pathway could be found in close proximity, either upstream or downstream, to the BPS cluster. The only unassigned *wza* gene in the chromosome was the orphan-like *mxan_1915* [17], separated from the BPS cluster by 1.015930 Mbp (**Fig 1B, S1C Fig**). Remarkably though, homology and synteny studies of related genomes revealed the gene encoding MXAN_1915 (now WzaB) to be contiguous with the genes encoding the other BPS assembly pathway members (i.e. WzcB, WzyB, and WzxB) in the genome of *Sandaracinus amylolyticus* (**Fig 1B, S1C Fig**), suggesting that MXAN_1915 may be functionally-linked with the BPS pathway.

Thus, the EPS, MASC, and BPS clusters all encode respective Wzx, Wzy, Wzc, and Wza proteins, despite the presence of large insertions between certain genes in the *M. xanthus* chromosome. While WzcB from the BPS pathway is a PCP-2A protein (i.e. contains a fused Wze-like C-terminal cytoplasmic BY-kinase domain), both WzcX and WzcS (from the EPS and MASC pathways, respectively) lack such a fusion and are PCP-2B proteins; instead, the EPS and MASC pathways encode stand-alone BY-kinase proteins (WzeX and WzeS, respectively) (**Fig 1A**).

### EPS and BPS directly affect community organization and behaviour

To better understand the functions of the EPS, BPS, and MASC assembly pathways, deletion-mutant strains were constructed for each pathway (**S1 Table**). In agreement with a previous report [44], all EPS-pathway mutants were severely compromised for swarm expansion and fruiting body formation (**Fig 2A,B**), confirming the central role of EPS in the *M. xanthus* life cycle. Additionally, MASC^−^ cells displayed WT-like group motility and fruiting body formation (**Fig 2A,B**), consistent with MASC pathway expression only during developmental phases [52]. To limit the scope of this project to vegetative cell physiology, minimal additional characterization of the MASC pathway is reported herein.

**Figure 2.**
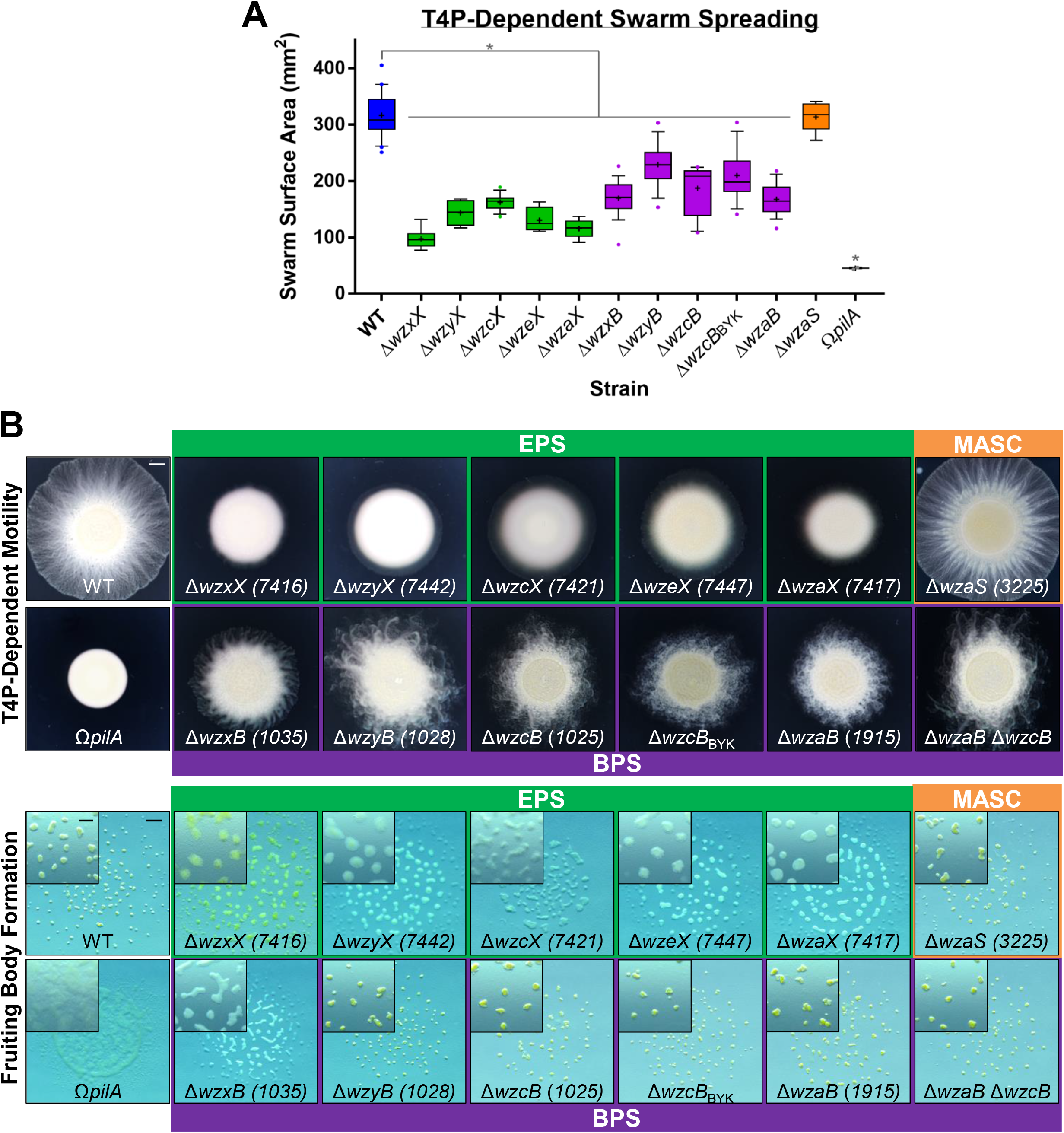
Physiological defects due to loss of EPS vs. BPS. **(A)** Box plots of the swarm surface obtained on 0.5% agar from T4P-dependent motility. The lower and upper boundaries of the boxes correspond to the 25^th^ and 75^th^ percentiles, respectively. The median (line through centre of boxplot) and mean (+) of each dataset are indicated. Lower and upper whiskers represent the 10^th^ and 90^th^ percentiles, respectively; data points above and below the whiskers are drawn as individual points. Asterisks denote datasets displaying statistically significant dataset differences (*p* < 0.05) compared to WT, as determined via 1-way ANOVA with Tukey’s multiple comparisons test. A minimum of 4 biological replicate values were obtained, each the mean of 3 technical replicates. Detailed statistical analysis are available (**S4 Table**). **(B)** EPS-, MASC-, and BPS-pathway mutant swarm physiologies. *Top:* T4P-dependent motility after 48 h (scale bar: 2 mm). *Bottom:* Fruiting body formation after 72 h (*main panel,* scale bar: 1 mm; *magnified inset,* scale bar: 400 µm).

Cells deleted for BPS-cluster genes displayed “fuzzy” swarm morphology during T4P-dependent swarm spreading (**Fig 2B**). Fruiting body formation in the absence of BPS also required lower cell densities (**Fig 2B, S2A Fig**), suggesting the BPS^−^ cells aggregate more easily. Neither BPS nor EPS was required for predation, as mutants defective in either pathway were still able to invade and digest a colony of prey *E. coli* cells (**S2B Fig**). Prey invasion is typically followed by the formation of ripples (i.e. synchronized waves of coordinated cells) (**S2B Fig**), a phenomenon that increases the efficiency of prey cell lysis [53]. While both BPS^−^ and EPS^−^ cells were still able to invade the *E. coli* colony, only BPS^−^ swarms displayed WT-like rippling, whereas EPS^−^ swarms did not ripple (**S2B Fig**). This suggests that (i) EPS may be required for predation, and if so (ii) BPS^−^ cells may still elaborate cell-surface EPS.

The only mutants that showed slightly divergent motility and developmental phenotypes compared to other respective EPS- and BPS-pathway mutants were Δ*wzxX* and Δ*wzxB* (**Fig 2**); this is consistent with *wzx* mutations in one pathway having the potential to affect the biosynthesis of polysaccharides from unrelated pathways (also requiring UndPP-linked precursors) due to depletion of available UndP [35, 54]. Importantly, the *mxan_1915* (now *wzaB*) mutant displayed identical phenotypes to those of the other BPS-pathway mutants (**Fig 2**), despite the 1.015930 Mbp separation of the gene from the rest of the BPS biosynthesis cluster (**Fig 1B**), further supporting the notion that *mxan_1915* indeed encodes for the BPS-pathway Wza (**Fig 1A**). In support of this hypothesis, a Δ*mxan_1915* Δ*wzcB* double mutant displayed phenotypes similar to those of the respective single mutants, consistent with *mxan_1915* (now *wzaB*) and *wzcB* encoding proteins that belong to the same (i.e. BPS) pathway (**Fig 2A,B**).

### BPS is not bound to the cell surface

The (i) hyper-aggregative character and (ii) rippling phenotype during predation of BPS^−^ cells (**S2 Fig**) suggested the presence of EPS on the surface of these cells. To address these observations, we set out to better understand the nature of the cell-surface EPS layer in BPS^−^ cells. As retention of the hydrophilic dye Trypan Blue has long been used as a readout for production of cell-surface EPS in *M. xanthus* [55], we obtained dye-binding profiles for all EPS- and BPS-pathway mutant strains. For all proposed EPS-pathway mutants, Trypan Blue binding was drastically reduced compared to WT cells (**Fig 3A**), consistent with previous descriptions of EPS deficiencies [55]. However, BPS-pathway mutants exhibited divergent Trypan Blue-binding profiles: (i) mutants unable to flip or polymerize UndPP-linked BPS repeat units (i.e. Δ*wzxB* and Δ*wzyB*) displayed significantly lower dye binding than WT cells (**Fig 3A**), consistent with reduced EPS production in these backgrounds. As discussed above, the likeliest explanation is that mutant strains in which UndPP-linked oligosaccharide repeat units for a particular pathway may build up can manifest polysaccharide synthesis defects in other pathways also requiring UndPP-linked units [54]. (ii) Conversely, *M. xanthus* BPS-pathway mutants with the capacity for periplasmic polymerization of the BPS chain, but compromised for BPS secretion (i.e. Δ*wzcB*, Δ*wzcB*_BYK_, Δ*wzaB*, and Δ*wzaB* Δ*wzcB*) did not display reduced Trypan Blue-binding relative to WT cells (**Fig 3A**). Compared to BPS-pathway Δ*wzaB*, the dye-binding profile of the EPS- and BPS-pathway double mutant Δ*wzaX* Δ*wzaB* matched that of the EPS-pathway Δ*wzaX* single-mutant (**Fig 3A**). Thus, the effect of BPS may be downstream to that of EPS.

**Figure 3.**
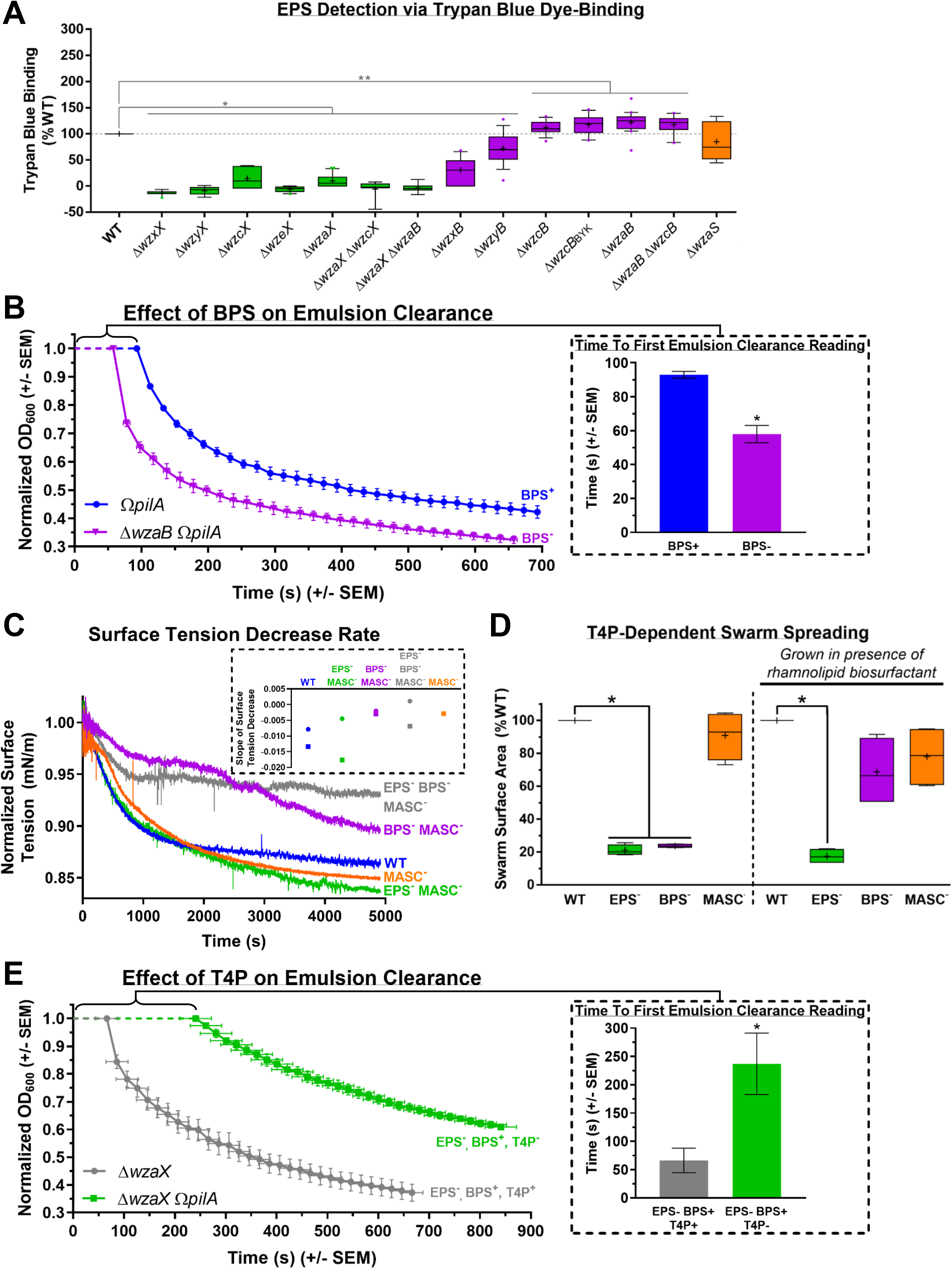
Analysis of BPS properties. **(A)** Boxplots of Trypan Blue dye retention to indicate the levels of EPS production in various strains relative to WT. The lower and upper boundaries of the boxes correspond to the 25^th^ and 75^th^ percentiles, respectively. The median (line through centre of boxplot) and mean (+) of each dataset are indicated. Lower and upper whiskers represent the 10^th^ and 90^th^ percentiles, respectively; data points above and below the whiskers are drawn as individual points. Asterisks denote datasets displaying statistically significant differences in distributions (*p* < 0.05) shifted higher (**) or lower (*) than WT, as determined via Wilcoxon signed-rank test performed relative to “100” (**S5 Table**). **(B)** Real-time clearance of hexadecane–CYE supernatant emulsions from BPS^+^ and BPS^−^ strains; values are the mean of 3 biological replicates (+/− SEM). OD_600_ values were normalized to their first registered values, whereas registered time points are displayed at their actual occurrence. *Inset:* Scanning time (post-mixing) for a given cuvette (containing a culture supernatant–hexadecane emulsion) until a first absolute value for OD_600_ could be registered by the spectrophotometer, for samples with(out) BPS (*n* = 3). Asterisk (*) denotes statistically-significant difference in mean value compared to *pilA* mutant (*p* = 0.0031), as determined via unpaired Student’s t-test. **(C)** Time course of normalized surface tension values (via digital drop tensiometry) from representative submerged-culture supernatants. Surface tension values across all time points were normalized against the initial surface tension value (*t* = 0) for each respective strain (**S3C Fig**). Strains tested: WT, MASC^−^ (Δ*wzaS*), BPS^−^ MASC^−^ (Δ*wzaB* Δ*wzaS*), EPS^−^ MASC^−^ (Δ*wzaX* Δ*wzaS*), EPS^−^ BPS^−^ MASC^−^ (Δ*wzaX* Δ*wzaB* Δ*wzaS*). *Inset:* Slope values from biological replicate time courses (each represented by a different shape) for each strain. Slopes were calculated by fitting the time-course curves with a fourth-degree polynomial function. **(D)** T4P-dependent swarm spreading in the presence of exogenous di-rhamnolipid-C_14_-C_14_ biosurfactant from *Burkholderia thailandensis* E264. The lower and upper boundaries of the boxes correspond to the 25^th^ and 75^th^ percentiles, respectively. The median (line through centre of boxplot) and mean (+) of each dataset are indicated. Lower and upper whiskers represent the 10^th^ and 90^th^ percentiles, respectively. Asterisks denote datasets displaying statistically significant differences in mean values (*p* < 0.05) compared to WT swarms, as determined via one-sample t-test performed relative to “100” (**S4 Table**). **(E)** Real-time clearance of hexadecane–CYE supernatant emulsions from T4P^+^ and T4P^−^ BPS-producing strains; values are the mean of 3 biological replicates (+/− SEM). OD_600_ values were normalized to their first registered values, whereas registered time points are displayed at their actual occurrence. *Inset:* Scanning time (post-mixing) for a given cuvette (containing a culture supernatant–hexadecane emulsion) until a first absolute value for OD_600_ could be registered by the spectrophotometer, for samples with(out) a functional T4P (*n* = 4). Asterisk (*) denotes statistically-significant difference in mean value compared to Δ*wzaX* (*p* = 0.0265), as determined via unpaired Student’s t-test.

Since the Trypan Blue-binding assay permits the detection of cell-associated polysaccharides as part of a contiguous surface matrix, the observation that dye binding by BPS^−^ cells was not reduced compared to WT cells suggests three possible scenarios: (i) BPS is surface-associated but does not bind Trypan Blue, (ii) BPS is instead secreted to the extracellular milieu, or (iii) the BPS-pathway machinery does not produce a polysaccharide and instead has a novel alternative function. The latter possibility is unlikely considering the abundance of putative GTase genes in the BPS cluster (**S3 Table**). The hypothesis that BPS is not cell-associated (and instead likely secreted) is supported by the results of monosaccharide analysis of cell-associated polymers from surface-grown WT and BPS^−^ submerged cultures; these analyses revealed that the major cell-associated sugar species are analogous in both composition and quantity between WT and BPS^−^ strains (**S3A Fig**), indicating that Trypan Blue is binding to the same polysaccharide target in each strain. Therefore, since WT and BPS^−^ cells elaborate comparable levels of EPS and do not display differences in surface-associated sugar content, we conclude that BPS does not remain attached to the cell surface.

### BPS is a secreted biosurfactant negatively-regulated by the T4P

Given the (i) aggregative phenotypes of BPS^−^ cells during *in vivo* fruiting body formation (**S2A Fig**), (ii) compromised swarm spreading on surfaces (**Fig 2A, B**), and (iii) lack of cell-associated sugar differences (**S3A Fig**), we reasoned that BPS may be a secreted surface-active biopolymer with emulsifying and/or surfactant properties that should therefore be found in the extracellular environment. We thus employed two independent methods to probe the surface-active properties of secreted BPS. The first approach takes advantage of the ability of both emulsifiers and surfactants to stabilize emulsions of two immiscible phases [56]. As *pilA* mutations reduce clumping of *M. xanthus* cells in liquid culture due to reduction (not elimination) of EPS production [57] (**S3B Fig**), these mutant strains can thus be grown to higher overall cell densities than piliated variants; we used this approach to naturally maximize the concentration of secreted polymers in our cultures. Enriched (aqueous) supernatants from dense cultures were thus extracted, filtered, and tested for the ability to stably mix with the hydrocarbon hexadecane [58, 59]; light transmission through samples in cuvettes was then monitored in real-time via spectrophotometry to monitor emulsion breaking. Whereas BPS^−^ supernatants demonstrated rapid phase separation, supernatants from BPS^+^ cultures formed more stable emulsions with hexadecane (**Fig 3B**); the latter samples even required 1.6× more time to simply allow any detectable light to pass through the sample from the start of attempted OD_600_ readings until the first registered value (**Fig 3B, *inset***). BPS thus possesses emulsion-stabilizing properties, a feature of both emulsifiers and surfactants.

In addition to increasing the stability of two immiscible phases (i.e. possessing emulsifying properties), to be considered a surfactant a given compound must also be able to reduce the surface/interfacial tension between two phases [56]. We thus also probed differences in surface tension between supernatants from submerged WT, EPS^−^, and BPS^−^ swarms using a highly-sensitive dynamic drop tensiometer. This method revealed a higher surface tension in BPS^−^ colony supernatants compared to supernatants from WT and EPS^−^ colonies that are still able to secrete BPS (**Fig 3C**). These data are consistent with surfactant properties of culture supernatants in strains with an intact BPS biosynthesis pathway.

To further test the function of BPS as a biosurfactant, we sought to rescue the motility phenotype of BPS^−^ swarms through addition of an exogenous biosurfactant, di-rhamnolipid-C_14_-C_14_ produced by *Burkholderia thailandensis* E264 [60]. This biosurfactant is not synthesized via a Wzx/Wzy-dependent polysaccharide assembly pathway; it is instead produced in the cytoplasm of its native bacterium requiring a 3-gene *rhl* cluster [60]. This renders rhamnolipid chemically distinct from high molecular weight biopolymers, while able to reduce surface tension through its strong surfactant properties. As reported, relative to WT swarms, EPS^−^ or BPS^−^ strains were severely compromised for T4P-dependent swarm spreading, while MASC^−^ swarms displayed no significant differences (**Fig 2A**). However, upon pretreatment of the agar surface with exogenous rhamnolipid biosurfactant, BPS^−^ swarms regained near-WT-like spreading (while EPS^−^ and MASC^−^ swarms displayed the same phenotypes as those in the absence of rhamnolipid) (**Fig 3D**). Exogenous biosurfactant addition is thus able to complement BPS deficiency *in trans*.

As EPS production has long been known to require the presence of a T4P [9, 22] (**S3B Fig**), we used the above-described hexadecane-based bioemulsifier assay and high-density cultures to test the T4P-dependence of surface-active BPS production. For supernatants from EPS-deficient BPS^+^ cultures encoding a T4P, gradually-breaking emulsions were observed for samples encoding a functional T4P; however, inactivation of the T4P in this parent strain gave rise to culture supernatants with remarkably-extended abilities to stabilize emulsions (**Fig 3E**). From the beginning of attempted OD_600_ readings until the first detected value, the latter samples took 3.6× as long just to allow any detectable light to pass through the cuvette (**Fig 3E, *inset***). Thus, as opposed to EPS production which requires a T4P [22], the presence of a T4P may have an inhibitory effect on the production of surface-active BPS.

### BPS is a randomly-acetylated repeating ManNAcA-rich tetrasaccharide

Finally, we set out to characterize the structure and composition of the novel secreted BPS molecule. Given the strong surface-active properties in enriched supernatants from BPS^+^ Δ*wzaX* Ω*pilA* cultures (**Fig 3E**), and lack thereof from BPS^−^ Δ*wzaB* Ω*pilA* enriched supernatants (**Fig 3B**), we studied the differences in polysaccharide content between these samples. Protein/nucleic acid-depleted samples were first analyzed via 1D proton NMR, revealing a cluster of peaks in the BPS^+^ enriched supernatant near 2 ppm that were not present in the BPS^−^ sample (**Fig 4A**). Samples were then separated via gel chromatography, revealing the presence of an acidic polysaccharide in the BPS^+^ sample (**S4A Fig**). Anion exchange chromatography followed by HSQC NMR analysis of the isolated polysaccharide revealed a complicated NMR spectrum due to random acetylation (**S4B Fig**). However, subsequent NMR analysis of a deacetylated sample revealed well-defined resonances (**Fig 4B, Table 1**). In the spectrum of the O-deacetylated polysaccharide, four spin systems were observed, each a pyranose sugar with a NHAc group at carbon 2, identified by ^13^C signal position between 50-60 ppm. All sugars displayed β-manno-configuration, as identified via TOCSY and NOESY signal patterns, and agreement with ^13^C signal positions (**Table 1**). One monosaccharide (residue A) contained a CH_2_OH group at carbon 6, and was thus designated *N*-acetyl-mannosamine. The remaining three sugars (residues B, C, and D) displayed no further correlations from hydrogen 5 and thus were classified as *N*-acetyl-mannosaminuronic acids, designations which were confirmed by the mass spectrum (**Fig 4C**). The sequence of the monosaccharides was identified based on NOE detection between nuclei A1:D3, B1:A4, C1:B4, D1:C4 (**Fig 4B**). Due to signal overlap, it was not possible to fully differentiate the latter two NOEs (**Fig 4B**), but this did not affect monosaccharide sequence designation.

**Figure 4.**
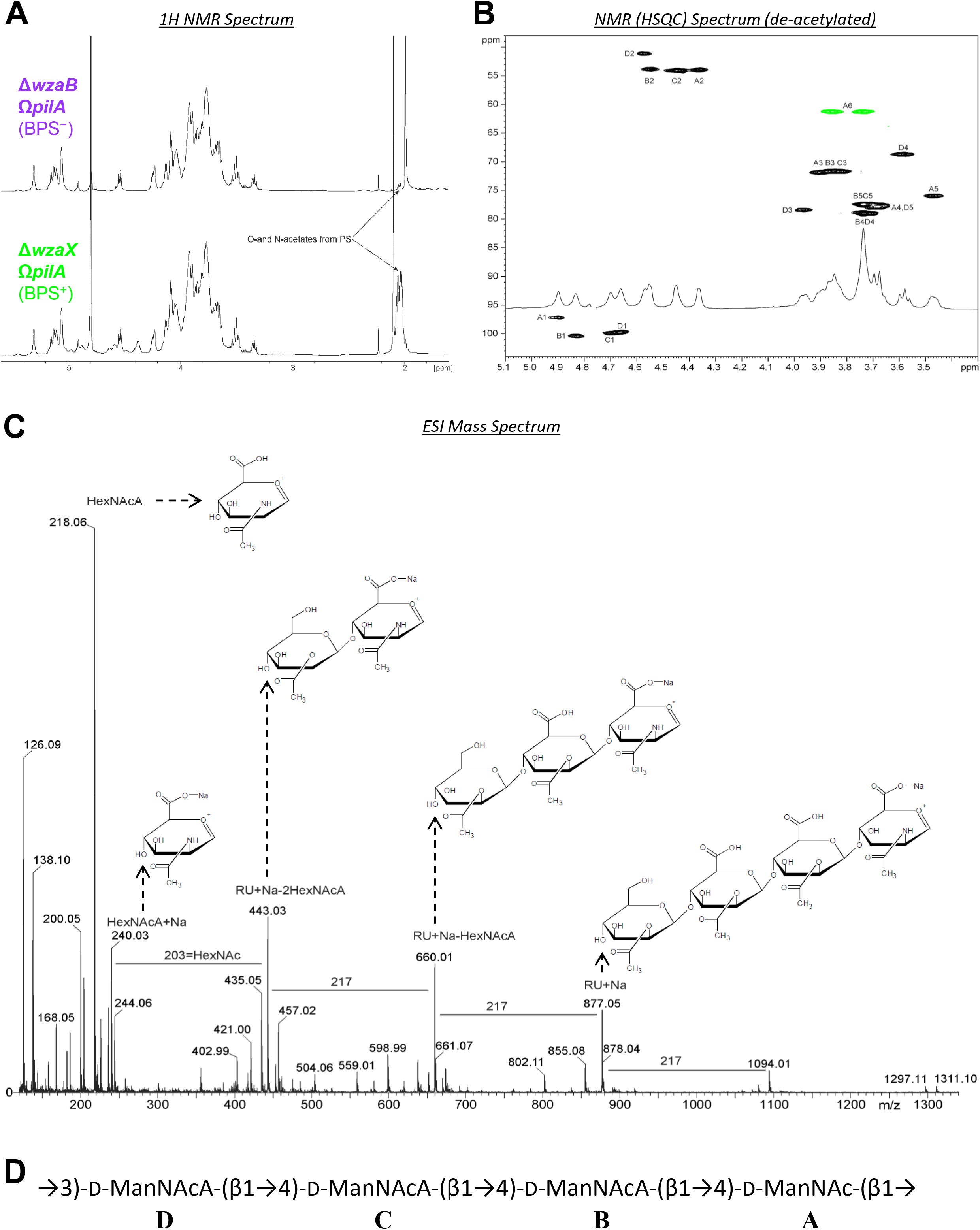
Analysis of BPS composition and structure. **(A)** 1H NMR spectra of concentrated supernatants from BPS^+^ Δ*wzaX* Ω*pilA* and BPS^−^ Δ*wzaB* Ω*pilA* cultures. **(B)** HSQC spectrum of deacetylated acidic polysaccharide isolated from Δ*wzaX* Ω*pilA* supernatant. Analysis was performed at 27 °C, 500 MHz. Resonance peak colours: *black*, C–H; *green*, C–H_2_. **(C)** Negative mode high cone voltage (180 V) ESI-MS of deacetylated acidic polysaccharide from Δ*wzaX* Ω*pilA* supernatant. RU = repeating unit. **(D)** Chemical structure of the BPS polymer tetrasaccharide repeating unit.

**Table 1.**
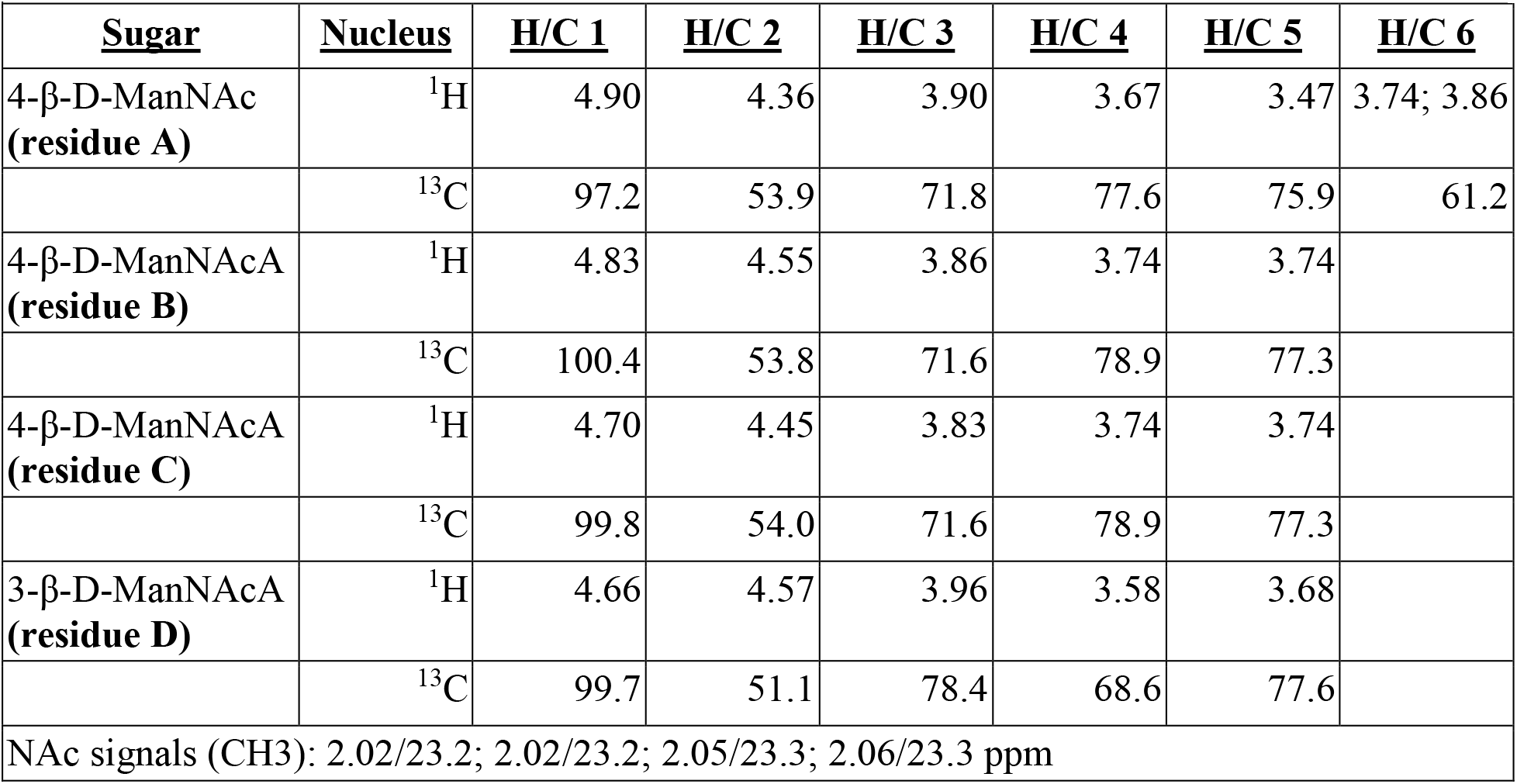
^1^H and ^13^C NMR data (δ, ppm, D_2_O, 27 °C, 500 MHz) for the deacetylated polysaccharide from Δ*wzaX* Ω*pilA* concentrated supernatant.

Using the information above, we were able to revisit the spectrum annotation for the untreated polysaccharide. The original polymer was found to be partially acetylated at position 3 in residues A, B, and C (**S4B Fig**). For residue D, oxygens 4 and 6 were capable of being acetylated, but no appropriate signal was visible; instead, positions 4 and 6 in residue D yielded the same signals as those detected in the deacetylated polysaccharide (**Fig 4B,C**). For residues A, B, and C, only oxygen 3 was free to be acetylated, thus agreeing with the signal position. Three OAc signals were present in the HSQC spectrum: 2.02/21.6; 2.04/21.6; 2.10/21.6 ppm, although there may be overlaps (**S4B Fig**).

Taken together, these analyses revealed BPS to be a heteropolysaccharide built of the following repeating unit: →3)-D-ManNAcA-(β1→4)-D-ManNAcA-(β1→4)-D-ManNAcA-(β1→4)-D-ManNAc-(β1→ (**Fig 4D**). Each tetrasaccharide repeat is thus joined via β1→3-linkages to another, with each repeating unit composed of a proximal neutral *N*-acetyl-β-D-mannosamine sugar, followed by three distal charged *N*-acetyl-β-D-mannosaminuronic acid sugars; a random acetylation pattern at position 3 was detected for the first three residues of the tetrasaccharide (**Fig 4D**). Therefore, the product of the Wzx/Wzy-dependent BPS assembly pathway (**Fig 1A**) is indeed a high molecular-weight biosurfactant polysaccharide (BPS) built of randomly-acetylated mannosaminuronic acid-rich tetrasaccharide repeat units (**Fig 4D**).

### BPS and EPS polymers are shared differently within the swarm community

Given the compromised nature of T4P-dependent swarm spreading for both EPS^−^ and BPS^−^ strains (**Fig 2**), we mixed populations of both strains together at different ratios to test the ability of each polysaccharide to cross-complement deficiency in the other and restore swarm motility on soft agar. Mixture of the two strains at a 1:1 ratio restored T4P-dependent swarm expansion to wild-type levels (**Fig 5A,B**), indicating an *in trans*-complementation of a motility defect. We next explored the spatial distribution of the two strains at a 1:1 ratio via fluorescence microscopy of EPS^−^ cells with a sfGFP-labelled OM and BPS^−^ cells elaborating a mCherry-labelled IM[18] (**Fig 5C**). While both EPS-pathway (*green*) and BPS-pathway (*red*) mutant cells were detected in the swarm interior as well as at its edge, the distribution was not homogeneous (**Fig 5C**); BPS^−^ cells (*red*, i.e. cells still able to produce EPS) were more abundant towards the centre of the swarm, whereas EPS^−^ cells (*green*, i.e. cells still able to produce BPS) were enriched toward the periphery (**Fig 5C**). At this ratio, the EPS^−^ cells in this mixture were thus able to utilize EPS produced by the BPS^−^ cells, indicating that EPS can be considered a “shared good” of the community. On the other hand, BPS^−^ cells remained at the swarm centre, suggesting that any BPS produced by the EPS^−^ cells was not able to rescue the motility defect of the BPS^−^ cells, suggesting that BPS may not be a “shared good”.

**Figure 5.**
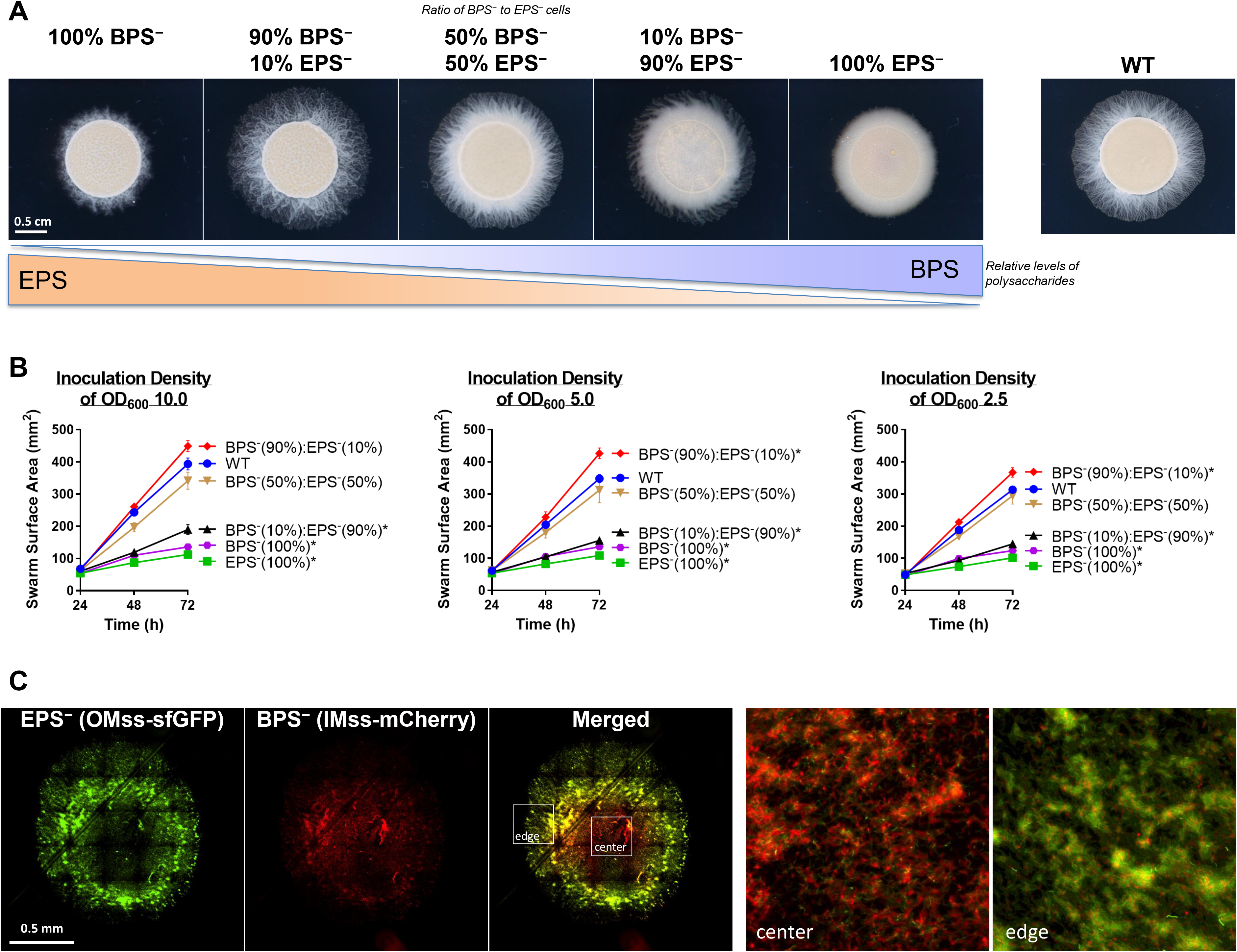
Cross-complementation of EPS vs. BPS deficiencies via strain mixing. **(A)** EPS^−^ (Δ*wzaX*) and BPS^−^ (Δ*wzaB*) cells from exponentially growing cultures were mixed at the indicated ratios to a final concentration of OD_600_ 10. Pure and mixed cultures were then spotted on CYE 0.5% agar and imaged after 48 h at 32 °C. **(B)** Swarm areas with temporal tracking of pure and mixed cultures, were treated as described and imaged at 24, 48, and 72 h. Each data point is the average of four biological replicates and is displayed +/- SEM. Mixed/pure cultures with statistically-significant differences (*p* ≤ 0.05) in mean surface areas (at 72 h relative to WT) are indicated with an asterisk (*), as determined via 1-way ANOVA followed by Dunnett’s multiple comparisons test. Detailed statistical analysis are available (**S4 Table**). **(C)** EPS^−^ (Δ*wzaX*) P*_pilA_*-OMss-sfGFP and BPS^−^ (Δ*wzaB*) P*_pilA_*-IMss-mCherry cells were mixed at a 1:1 ratio as in (A), spotted on agar pads, and imaged via fluorescence microscopy after 24 h. The two images on the right are magnified views of the colony centre and colony edge approximately indicated by the inset boxes in the “merged” image.

### Expression of BPS vs. EPS is spatially and temporally distinct within a swarm

Considering the distinct physiologies of EPS^−^ vs. BPS^−^ cells and swarms (**Fig 2, 3, S2,3 Fig**), we set out to examine whether or not EPS and BPS were differentially regulated by cells in WT swarm communities. To first probe the spatial and temporal distribution of EPS- and BPS-pathway cluster expression within a swarm, sfGFP and mCherry were simultaneously placed under EPS- and BPS-cluster promoter control (P_EPS_-sfGFP [*wzxX* promoter] and P_BPS_-mCherry [*wzcB* promoter], respectively) in the same WT cell. The spatiotemporal expression of the two reporter genes was subsequently monitored within the swarm via a recently-developed fluorescence microscopy technique allowing for the acquisition of large-scale images at high resolution [61]. While cells plated immediately from liquid cultures (*t* = 0) demonstrated homogeneous expression profiles of both reporters across the population, spatial separation of signals within a swarm was observed already after 24 h (**S5 Fig**). These distinct spatial signal profiles were even more pronounced after 48 h, with the P_BPS_-mCherry signal preferentially expressed towards the swarm interior, whereas P_EPS_-sfGFP signal was more highly expressed around the swarm periphery (**Fig 6A**). Fruiting bodies that appeared after 72 h indicated signal overlap between P_EPS_-sfGFP and P_BPS_-mCherry expression, suggesting that EPS and BPS pathways are both active within these multicellular structures; at this time point, P_EPS_-sfGFP expression was still detectable at the swarm edge (**S5 Fig**).

**Figure 6:**
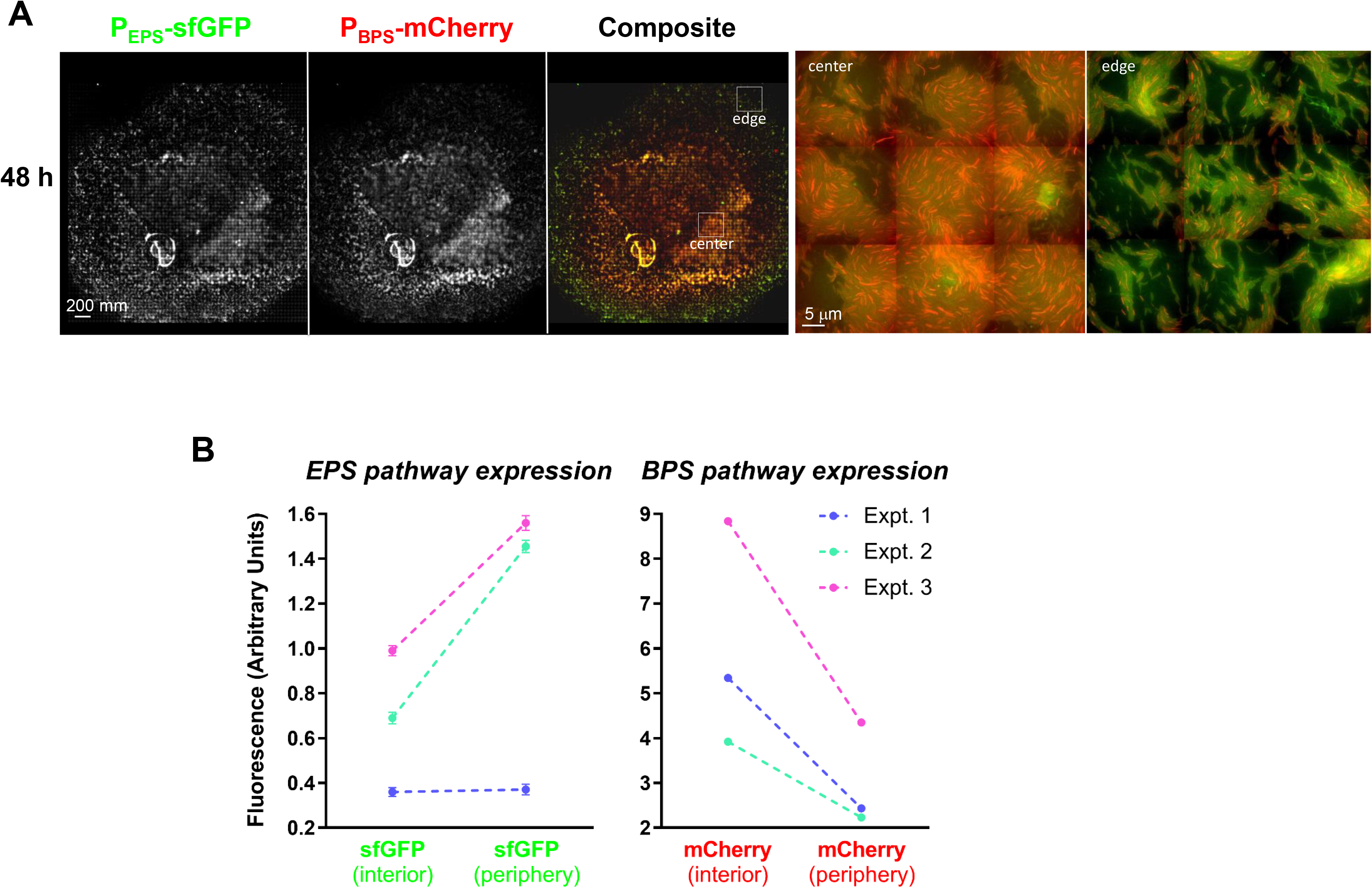
Analysis of the spatial expression of the EPS and BPS gene clusters. **(A)** Dual-labelled (P_EPS_-sfGFP + P_BPS_-mCherry) WT cells (strain EM709) from exponentially growing cultures were spotted on developmental media at a final concentration of OD_600_ 10.0 and imaged at 48 h. Images were scaled as described in Material and Methods. The two images on the right are magnified views of the colony centre and colony edge at positions approximately indicated by the inset boxes in the composite image. **(B)** FACS analysis of WT + P_EPS_-sfGFP + P_BPS_-mCherry cells collected from the colony centre (int.) or edge (ext.) following incubation for 48 h. Cells were analyzed for intensity of sfGFP and mCherry fluorescence. Results of three independent experiments are displayed (1^st^ experiment, *orange* lines; 2^nd^ experiment, *blue* lines; 3^rd^ experiment, *magenta* lines). For each experiment, a total population of 300,000-500,000 events was used and statistically analyzed. Differences between fluorescence intensity at the colony center vs. edges are significant (*p* < 0.0001) for all experiment except for the first experiment with GFP. Errors bars are set at 1% confidence. Signals obtained with the non-fluorescent wild-type strain were subtracted from the fluorescence signals of strain EM709.

To quantify the differential P_EPS_-sfGFP and P_BPS_-mCherry expression patterns observed via fluorescence microscopy (**Fig 6A**), we collected cell samples from the edges and the centres of swarms after 48 h and analyzed these fractions via fluorescence-activated cell sorting (FACS). While red fluorescence (from P_BPS_-Cherry) was 2× more intense towards the colony interior compared to the periphery, the inverse relationship was observed for green fluorescence; P_EPS_-sfGFP expression was 2× more intense at the periphery versus the swarm centre (**Fig 6B**). These FACS data directly reinforce the spatial expression patterns described above (**Fig 6A**).

All together these data indicate that the production of EPS and BPS occurs with a precise spatiotemporal regulation likely reflecting the requirement for the two polysaccharides in specific cell behaviours within different swarm regions. For example, EPS may be preferentially required at the colony edge where T4P-dependent swarm spreading takes place, whereas BPS is more crucial at the high cell-density colony centre to perhaps favour initial cell dispersal.

## Discussion

Given the extensive genomic, phenotypic, biochemical, and biophysical characterizations described above, we propose that the product of the BPS assembly pathway is a novel T4P-regulated acidic heteropolysaccharide with surface-active properties. These characteristics enhance swarm expansion. To our knowledge, this is the first report of a product synthesized by a Wzx/Wzy-dependent polysaccharide assembly pathway with surfactant properties. Importantly, these data directly affect past, present, and future interpretations of *M. xanthus* physiology attributed to effects on EPS, which could now be due in part to the differential regulation of BPS.

In this investigation, we have demonstrated that *M. xanthus* uses temporally- and spatially-distinct polysaccharide production patterns amongst members of the swarm community to modulate its complex multicellular lifecycle, with differently-localized subpopulations of cells favoring production of EPS vs. BPS to promote dissemination of the swarm community. This dynamic thus facilitates the development of complex swarm-level behaviours requiring more than the activity of a collection of single cells. It is in this manner that the collective behaviour of *M. xanthus* cells in swarms leads to the differentiation of the population into forager cells, peripheral rods, and spore-forming cells.

The combined importance of secreted polymers and surface-active agents for community expansion and migration may be a common theme for bacteria, especially for species that display differentiated cell fates. In the model Gram-positive bacterium *Bacillus subtilis*, at least five cell types have been described to date, with each distinguished via the following phenotypes: protease production, sporulation, motility, matrix production, and surfactin production [62, 63]. (i) Extracellular protease production is correlated with later stages of biofilm formation, suggesting a role in nutrient acquisition or escape from a biofilm [64]. (ii) Sporulation allows for *B. subtilis* to survive prolonged periods of desiccation and nutrient scarcity, whereas (iii) motility phases involve synthesis of external appendages (flagella) for swimming [63]. As with numerous other bacteria, (iv) *B. subtilis* also biosynthesizes an extracellular polysaccharide that constitutes the bulk of its biofilm matrix, and which under certain conditions can affect colony expansion. (v) Finally, *B. subtilis* also famously produces surfactin, a cyclic lipopeptide with strong surfactant properties [65–67]. Recently, the formation of matrix polysaccharide-dependent “van Gogh” bundles of *B. subtilis* cells at the colony edge was shown to be greatly improved in the presence of nearby surfactin-producing cells; the net effect of this interplay between matrix polysaccharide and surfactin was thus to determine the rate of colony expansion [6]. Intriguingly, each of the differentiating phenotypes describe above has a direct parallel in *M. xanthus*. (i) Numerous proteases are released by *M. xanthus* cells and are suspected to participate in nutrient acquisition (via predation) [68], (ii) whereas myxospore formation in fruiting bodies promotes survival under nutrient-limiting conditions [13]. Though single-cell gliding motility does not require any external appendages [12], (iii) swarm-level motility is mediated by extension and retraction of T4P appendages; (iv) this is in conjunction with the requirement of EPS for T4P-dependent swarm spreading [13]. (v) As reported herein, *M. xanthus* also produces a surface-active BPS molecule, in the form of a high molecular-weight heteropolysaccharide polymer, with the effect of BPS on EPS modulating swarm-level ultrastructure, migration, and the overall developmental cycle of the bacterium.

The production of a secreted polysaccharide with emulsifying properties and its interaction with cell-surface-associated carbohydrates also has parallels in another well-studied system. Originally isolated from a mixed culture growing on crude oil [69], *Acinetobacter* sp. RAG-1 was found to produce a compound able to stably emulsify hydrocarbons [70, 71]. Termed RAG-1 emulsan, this compound has been extensively tested with respect to environmental applications including oil spill cleanup and bioremediation of heavy metal pollution [72, 73]. RAG-1 emulsan had been studied for over 35 years [69] before a more rigorous extraction protocol was developed, revealing that RAG-1 emulsan was in fact a bipartite compound composed of (i) a rough-type LPS and (ii) a high molecular-weight secreted polysaccharide [74]. The latter compound, now termed *Acinetobacter* Polyelectrolytic Exopolysaccharide (APE)[75], is synthesized via a Wzx/Wzy-dependent pathway [76, 77] and is the component responsible for the emulsifying properties of RAG-1 emulsan [74]. To our knowledge, the capacity of APE to reduce interfacial tension has never been reported; thus its role as a true biosurfactant has not been established. As the APE polymer is built from repeating →4)-D-GalNAc-6-OAc-(α1→4)-D-GalNAcA-(α1→3)-D-QuiNAc4NHb-(α1→ trisaccharide units [75], it is conceivable that the presence of uronic acid and acetyl moieties are able to contribute hydrophilic and hydrophobic character to APE, respectively. Since *M. xanthus* BPS is composed of →3)-D-ManNAcA-(β1→4)-D-ManNAcA-(β1→4)-D-ManNAcA-(β1→4)-D-ManNAc-(β1→ repeating units, considerable hydrophilic and hydrophobic character would again be present due to its high uronic acid content as well as its N-/O-acetylation levels, respectively. Together, these traits would explain the emulsifying properties of both APE and BPS (as well as the surface tension-reducing surfactant properties in the latter). Similarly to *Acinetobacter* and RAG-1 emulsan, physiology connected directly to the presence/absence of “EPS” in *M. xanthus* has been studied for over 30 years [78]; however, the complex physiology of this social bacterium must now be considered within the context of the interplay between the dedicated EPS polymer on the cell surface and the newly-identified secreted BPS product.

With the identification of BPS and its importance to *M. xanthus* physiology, numerous questions have been raised. For instance, given the spatiotemporal differences in production of the various polymers, by what mechanism is BPS production regulated in relation to EPS and MASC? The presence of functionally-important BY-kinase homologues to the production of BPS/EPS/MASC represents an enticing level of production control for each of the polymers. In addition, the apparent suppression of BPS production by the presence of a T4P suggests possible links between BPS and the Dif pathway, with the latter already known to regulate T4P-dependent EPS expression. As biosurfactants are of immense industrial interest and importance, further characterization of the chemical properties of BPS are also immediate avenues of future inquiry.

Ultimately, our investigation reveals that differentiated functions between distinct cell subpopulations across an entire swarm generates an ecologically-beneficial, community-level, higher-order organization. This is a central tenet governing the varied evolutionary origins of multicellularity in nature.

## Materials and Methods

### Bacterial Cell Culture

The *M. xanthus* strains used in this study are listed in **S1 Table**. They were grown and maintained at 32 °C on Casitone-yeast extract (CYE) agar plates or in CYE liquid medium at 32°C on a rotary shaker at 160 rpm. The *Escherichia coli* strains used for plasmid construction were grown and maintained at 37 °C on LB agar plates or in LB liquid medium. Plates contained 1.5% agar (BD Difco). Kanamycin at 100 μg/mL and galactose at 2.5% (w/v) was added to media for selection when appropriate.

### Plasmid and Mutant Construction

Plasmids used in this study are listed in **S1 Table**. To create *M. xanthus* in-frame deletion strains, 900 bp upstream and downstream of the gene targeted for deletion were amplified and fused via PCR, restriction digested, then ligated into pBJ113 or pBJ114 [79]. The resulting plasmids were then introduced into wild-type *M. xanthus* DZ2 via electroporation. Mutants resulting from homologous recombination of the deletion alleles were obtained by selection on CYE agar plates first containing kanamycin, and then containing galactose to resolve the merodiploids.

To fluorescently label the IM or outer membrane (OM) of the *M. xanthus* EPS^−^ (Δ*wzaX*) and BPS^−^ (Δ*wzaB*) strains, cells were electroporated with the integrative vector pSWU19 encoding either the OMss-sfGFP or the IMss-mCherry fusion, respectively, under control of the *pilA* promoter [18]. Strain EM709 was obtained by amplifying 1000 bp upstream of the start codon of genes *mxan_7416* and *mxan_1025* (https://www.genome.jp/kegg/) and fusing them to the *sfgfp* or *mcherry* coding sequences, respectively. The two gene fusions were then cloned into the same pSWU19 vector. The recombinant vector was then transferred via electroporation into *M. xanthus* DZ2, thus inserting the two gene fusions together at the Mx8 phage-attachment site in the chromosome [18] and allowing for co-expression in *M. xanthus*.

### Phylogeny and Gene Co-Occurrence

Forty-order Myxococcales genomes[80–95] were downloaded from NCBI followed by RAST-based gene prediction and annotation[96]. Thirty housekeeping proteins (DnaG, Frr, InfC, NusA, Pgk, PyrG, RplC, RplD, RplE, RplF, RplK, RplL, RplM, RplN, RplP, RplS, RplT, RpmA, RpoB, RpsB, RpsC, RpsE, RpsI, RpsJ, RpsK, RpsM, RpsS, SmpB, Tsf) were aligned, concatenated, and subjected to FastTree 2.1.8 to generate a maximum likelihood phylogeny with 100 bootstrap values using the Jones–Taylor–Thornton (JTT) protein substitution model. Functional domains were identified by scanning all proteomes against the Pfam-A v29.0 database [97] (downloaded: Oct 26, 2016) using hmmscan (E-value cutoff 1e-5) from the HMMER suite (http://hmmer.janelia.org/) [98] and further parsed using hmmscan-parser.sh. PFAM domains attributed to Wzx (MXAN_7416; Polysacc_synt), Wzy (MXAN_7442; PF04932 [Wzy]), Wzc (MXAN_7421; PF02706 [Wzz]), and Wza (MXAN_7417; Poly_export) were identified and the protein information was extracted. Based on the identified proteins and their location in the genome, clusters were manually curated. Along with Pfam domain analysis, we used all protein sequences forming identified clusters to perform Basic Local Alignment Search Tool (BLASTp) searches [99] against the predicted proteome of each organism using stringent cut-offs [E-value of 0.00001, 35% query coverage and 35% similarity]. To confirm the participation of identified homologs (via PFAM and BLAST) in respective clusters, we also generated maximum likelihood phylogenetic trees (JTT Model and 100 bootstrap values using FastTree 2.1.8) of homologs MXAN_7416, MXAN_7417, MXAN_7421, and MXAN_7442. Finally, the binary distribution (Presence/absence), location and cluster information of each cluster was mapped on the housekeeping protein-based phylogeny. Consensus α-helical transmembrane segment (TMS) prediction [100] was obtained via OCTOPUS[101] and TMHMM[102]. Fold-recognition analyses were performed via HHpred [103].

### Phenotypic Analysis

Exponentially-growing cells were harvested and resuspended in TPM buffer (10 mM Tris-HCl, pH 7.6, 8 mM MgSO_4_ and 1 mM KH_2_PO_4_) at the final concentration of OD_600_ 5.0 or swarming assays or OD_600_ 1.5, 2.5, 5.0, and 10.0 for developmental assays. This cell suspension (10 μL) was spotted onto CYE 0.5% agar or CF 1.5% agar for swarming or developmental (i.e. fruiting body formation) assays, respectively. Plates were incubated at 32 °C for 48 h (swarming) or 72 h (fruiting body formation) and photographed with an Olympus SZ61 binocular stereoscope. To examine the effect of exogenous biosurfactant on swarming, 100 µL of *Burkholderia thailandensis* E264 di-rhamnolipid-C_14_-C_14_ stock solution (300 ppm, 0.2 µm^2^-filtered) was first spread on top of the agar surface and allowed to dry in a biohood prior to inoculation. Swarm contours in each image were defined using cellSens software (Olympus), followed by calculation of surface area.

For strain-mixing time course experiments, 10 µL of TPM resuspensions at OD_600_ 2.5, 5.0, and 10.0 were spotted on square (15 cm × 15 cm) CYE 0.5% agar plates, with all six samples for each particular OD_600_ spotted on the same plate. Four biological replicates were analyzed for each set of parameters. Plates were inverted and incubated at 32 °C, with swarm images captured at 24, 48, and 72 h time points using an Olympus SZX16 stereoscope with UC90 4K camera. Swarm contours in each image were defined using the cellSens software suite (Olympus), followed by calculation of surface area via pixel quantitation.

### Fluorescence Microscopy

For the imaging of colonies by fluorescence microscopy, samples were prepared as described by Panigrahi and colleagues [61]: exponentially-growing cells were harvested and resuspended as pure or mixed culture in TPM buffer at the final concentration of OD_600_ 5.0. This cell suspension (1 μL) was spotted onto thin pads of CYE 1% agar for swarming assays and CF 1.5% agar for developmental and predation assays. For the latter, 1 μL of exponentially-growing *E. coli* MG1655 was also added (resuspended to OD_600_ 5.0). To avoid the desiccation of the thin agar pads, the agar was poured onto squared adhesive frames previously pasted on to glass slides. Slides were then incubated at 32 °C for 0, 24, 48 and 72 h, covered with the coverslips and photographed with a Nikon Eclipse TE2000 E PFS inverted epifluorescence microscope. Slides were imaged with a 10× objective for the strain-mixing experiments, and with a 100× objective for imaging of the dual-labelled strain. For each slide, a series of images was automatically captured by the aid of the Nikon Imaging Software to cover a section of the swarm via tiling and stitching. The microscope devices were optimized in order to minimize the mechanical movement and provide rapid autofocus capability (epi/diascopic diode lightening, piezo-electric stage) as previously described [79]. The microscope and devices were driven by the Nikon-NIS “JOBS” software [79].

### Trypan Blue Dye Retention

Trypan Blue dye-retention analysis was adapted from a previous report [55]. Cells from overnight cultures were sedimented and resuspended in TPM buffer to OD_600_ 1.0, after which 900 µL of cell resuspension was transferred to a 1.5 mL microfuge tube; a cell-free blank was also prepared with an identical volume of TPM. To each tube, 100 µL of Trypan Blue stock solution (100 µg/mL) was added, followed by a brief 1 s pulse via vortex to mix the samples. All tubes were placed in a rack, covered with aluminum foil, and incubated at room temperature on a rocker platform for 1 h. After this dye-binding period, samples were sedimented at high speed in a tabletop microfuge (16 000 × *g*, 5 min) to clear supernatants of intact cells. So as not to disrupt the pellet, only 900 µL of the dye-containing supernatant was aspirated and transferred to a disposable cuvette. The spectrophotometer was directly blanked at 585 nm (the absorption peak for Trypan Blue) using the cell-free “TPM + Trypan Blue” sample to calibrate the baseline absorbance corresponding to no retention of Trypan Blue dye by cells. Absorbance values at 585 nm (A_585_) were obtained for each clarified supernatant and normalized as a percentage of the WT A_585_ reading (i) as an internal control for each individual experiment, and (ii) to facilitate comparison of datasets across multiple biological replicates. Negative final values are due to trace amounts of cell debris detected at 585 nm in individual samples in which absolutely no binding of Trypan Blue occurred.

### Purification and Monosaccharide Analysis of Cell-Associated Sugars

*M. xanthus* cell-associated sugars were purified from CYE 0.5% agar-grown cultures as described [104], with the following modifications. Cell cultures were harvested and resuspended in 25 mL TNE buffer (100 mM Tris pH 7.5, 100 mM NaCl, 5 mM EDTA). Cells in suspension were then disrupted via sonication (4 pulses, 15 s each), followed by addition of SDS to a final concentration of 0.1% to extract cell-associated sugars. To remove DNA from cell samples, lysates were treated with 20 µL of 10 kU DNase I from LG Healthcare (20 mM Tris-HCl pH 7.6, 1 mM MgCl_2_, 50% v/v glycerol) and incubated at room temperature for 5 min. To obtain protein-free extracellular sugar samples, 1 mg/mL of Pronase E protease mixture (Sigma–Aldrich) was gently added directly and allowed to incubate at 37 °C (2 h). The extracts were sedimented (10 min, 7513 × *g*, 17 °C) and the pellets were washed twice with 25 mL TNE+SDS. The pellets were then washed twice with 10 mL TNE to remove any remaining SDS. To remove EDTA, sugar samples were washed twice with MOPS buffer (10 mM MOPS [pH 7.6], 2 mM MgSO_4_) and twice with cohesion buffer (10 mM MOPS buffer [pH 6.8], 1 mM CaCl_2_ 1 mM MgCl_2_). Finally, sugar samples were stored in cohesion buffer at −80 °C until use.

Cell-associated sugars purified from colonies (50 µL), were mixed with 500 µL of 12 M H_2_SO_4_ and incubated for 1 h at 37 °C under mild shaking; 20 µL of each sample were then mixed with 220 µL of distilled water, and the diluted samples were autoclaved for 1 h at 120 °C. After cooling, 50 µL of 10 M NaOH were added, and the samples were sedimented (10 000 × *g*, 10 min, room temperature) [105]. Supernatant (5 µL) was mixed with ddH_2_O (245 µL), followed by identification and quantification of the released monosaccharides via high-performance anion-exchange chromatography coupled with pulsed amperometric detection (HPAEC-PAD), performed in a Dionex ICS 3000 (Thermo Scientific) equipped with a pulsed amperometric detector. Sugar standards or EPS hydrolysates (25 µL) were applied to a Dionex CarboPac PA20 column (3 × 150 mm) preceded by the corresponding guard column (3 × 30 mm) at 35 °C. Sugars were eluted at 0.45 mL/min with the buffers 0.1 M NaOH, 1 M sodium acetate + 0.1 M NaOH and ddH_2_O as the eluents A, B and C, respectively. The following multi-step procedure was used: isochratic separation (10 min, 27% A + 73% C), gradient separation (20 min, 2 – 19 % B + 98 – 81% C), column wash (5 min, 100% A) and subsequent column equilibration (10 min, 27% A + 73% C). Injection of samples containing glucose, rhamnose, *N*-acetylglucosamine, arabinose, xylose, mannose, galactose, and glucosamine (Sigma-Aldrich) at known concentrations (ranging from 5 to 100 µM) were used to identify and quantify the released monosaccharides.

### Purification and Analysis of Secreted Polysaccharides

Lyophilized concentrated samples representing 175 mL of original culture supernatant for each strain were resuspended in ddH_2_O and treated with 2% acetic acid (10 min, 80 °C) to precipitate proteins and nucleic acids. Solutions were then separated via gel chromatography on a Sephadex G-15 column (1.5 cm × 60 cm) or Biogel P6 column (2.5 cm × 60 cm), in 1% acetic acid, monitored by refractive index detector (Gilson).

Anion-exchange chromatography was then performed via sample injection into a HiTrapQ column (Amersham, two columns × 5 mL each, connected together) in ddH_2_O at 3 mL/min. Samples were washed with ddH_2_O for 5 min, then eluted with a linear gradient from ddH_2_O to 1 M NaCl over 1 h with UV detection at 220 nm. Spot tests were performed on silica TLC plates, developed by dipping in 5% H_2_SO_4_ in ethanol and heating with heat gun until brown spots became visible. Samples were desalted on a Sephadex G-15 column.

NMR experiments were carried out on a Varian INOVA 500 MHz (^1^H) spectrometer with 3 mm Z-gradient probe with acetone internal reference (2.225 ppm for ^1^H and 31.45 ppm for ^13^C) using standard pulse sequences for gCOSY, TOCSY (mixing time 120 ms), ROESY (mixing time 500 ms), and gHSQCAD. Resolution was kept <3 Hz/pt in F2 in proton-proton correlations and <5 Hz/pt in F2 of H-C correlations. The spectra were processed and analyzed using the Bruker Topspin 2.1 program.

Monosaccharides were identified by COSY, TOCSY and NOESY cross peak patterns and ^13^C NMR chemical shifts. Aminogroup location was concluded from the high-field signal position of aminated carbons (CH at 45-60 ppm).

Electrospray-ionization (ESI) mass spectrometry (MS) was performed using a Waters SQ Detector 2 instrument. Samples were injected in 50% MeCN with 0.1 % TFA.0

### Emulsification Testing

Overnight *M. xanthus* cultures (50 mL CYE in 250 mL flasks) were inoculated at an initial OD_600_ of 0.05 and grown at 32 °C with shaking (220 rpm) to saturation (OD_600_ ∼ 5.0-7.0). Cultures were transferred to a 50 mL conical tube and sedimented at 7000 × *g* (25 min, 22 °C, JA-17 rotor). Supernatants were decanted into a syringe, passed through a 0.22-micron filter to remove remaining cells, and transferred (4 mL) to a quartz cuvette, followed by addition of 300 µL hexadecane (Sigma) coloured with Sudan Black dye (0.1 g of Sudan Black powder per 50 mL of hexadecane). Each cell-free supernatant sample was vigorously mixed with the coloured hexadecane 250 times over 2 min (via aspiration/ejection with a p1000 micropipette). Cuvettes were then immediately inserted into a spectrophotometer, with continual, rapid manual attempts made to obtain an initial OD_600_ reading, with this time recorded. After obtaining an initial OD_600_ reading, subsequent OD_600_ readings were manually carried out at 20 s intervals over 10 min to monitor the rate of emulsion clearance. All OD_600_ readings for each time course were normalized with respect to the initial OD_600_ value detected for each sample.

### Surface Tension Testing

The adsorption and interfacial properties as a function of time for supernatants of the five strains (and their secreted polysaccharides) were analyzed by means of a digital Tracker Drop Tensiometer (Teclis, Civrieux-d’Azergues, France) [106] at room temperature. From digital analysis of a liquid drop or an air bubble profile collected by a high-speed CCD camera, characteristic parameters (surface tension, area, volume) were determined in real time. Surface tension was estimated from the Laplace equation adapted for a bubble/drop. By controlled movements of the syringe piston, driven by a step-by-step motor, surface area can be maintained constant during the whole experiment. Before each experiment, cleanliness of material was tested using ultrapure water, before being dried with argon. For the study of tensioactive properties of the supernatants, a 10 µL air bubble was formed at the tip of a J-tube submerged in 5 mL of supernatants for each strain.

### Flow Cytometry

The fluorescence intensity of *M. xanthus* strain EM709 simultaneously expressing P_EPS_-sfGFP and P_BPS_-mCherry were measured by Fluorescence-Activated Cell Sorting (FACS) with a Bio-Rad S3E cells sorter. The blue laser (488nm, 100mW) was used for the forward scatter (FSC), side scatter (SSC) and excitation of sfGFP, whereas the green laser (561nm, 100mW) for the excitation of mCherry. Signals were collected using the emission filters FL1 (525/30 nm) and FL3 (615/25 nm) for sfGFP and mCherry, respectively. Cells collected from the colony edges and centers were suspended in TPM and ran at low-pressure mode and at a rate of 10,000 particles/s. The threshold on FSC was 0.12 and the voltages of the photomultipliers were 361, 280, 785 and 862 volts for FSC, SSC, FL1 and FL3, respectively. The density plots obtained (small angle scattering FSC versus wide angle scattering SSC signals) were gated on the population of interest and filtered to remove multiple events. Populations of 300,000 to 500,000 events were used and analyzed statistically using the FlowJo software. The sfGFP and mCherry signals obtained with the non-fluorescent wild type cells were subtracted from the signals obtained with cells of strain EM709. Measurements were carried out three times with bacteria from different plates.

## Acknowledgements

The authors would like to thank several individuals: (i) Mariamichela Lanzilli for constructing strain EM709; (ii) Lucie Lacombe for acquisition optimization of large-scale fluorescence microscopy images; (iii) Jean-François Guillemot for writing the script permitting viewing of large-scale fluorescence microscopy images; (iv) Éric Déziel for insightful discussions, troubleshooting regarding emulsifier and hydrophobicity testing, providing rhamnolipid, as well as critical reading of the manuscript; (v) Alec McDermott for assistance with dye-binding assays; (vi) Philippe Constant for valuable input on biostatistics. A Discovery operating grant (RGPIN-2016-06637) from the Natural Sciences and Engineering Research Council of Canada and a Discovery Award (2018-1400) from the Banting Research Foundation fund work in the lab of S.T.I. as well as studentships for F.S. and N.Y.J.; the latter two are also recipients of graduate studentships from the PROTEO research network. S.T.I. was supported by a post-doctoral fellowship in T.M.’s group at project inception from the Canadian Institutes of Health Research and the AMIDEX excellence program of Aix-Marseille University. Research in the lab of E.M.F.M. is supported through the Agence National de la Recherche (ANR-14-CE11-0023-01). I.V. is supported by a studentship from the CONACYT of Mexico. Work from the M.S. lab was supported by a grant from the National Science Foundation (IOS135462). None of the abovementioned funding sources had any input in the preparation of this article, or in the work described herein.

## Author Contributions

STI and EMFM conceived of and planned the study.

STI, IVA, and FS performed stereoscopic phenotypic analyses.

AG, FS, and IVA generated mutant constructs and strains.

IVA and FS measured colony surface areas, with the latter performing rhamnolipid trans-complementation.

AG and FS performed swarm-mixing experiments.

EMFM and GB completed FACS analyses.

EMFM and CM carried out epifluorescence microscopy.

AG completed predation assays.

IVA and HPF analyzed cell-associated polysaccharides.

EV and GR analyzed supernatant-derived polysaccharides.

GS generated phylogenetic data, with analysis by STI and GS.

STI performed protein sequence and fold-recognition analyses.

STI and FS performed dye-binding and auto-aggregation assays.

FS tested emulsion clearance.

AB, JLB, IVA and FS tested surfactant properties.

STI and EMFM wrote the manuscript.

STI and EMFM generated figures.

STI, EMFM, HPF, MS, CG, and TM contributed personnel and/or funding support.

