## Supplementary Figures 1-5 for "Modulation of bacterial multicellularity via spatiotemporal polysaccharide secretion"

#### SUPPORTING INFORMATION FIGURE LEGENDS:

**S1 Figure.** Gene conservation and synteny diagrams for **(A)** EPS, **(B)** MASC, and **(C)** BPS cluster evolution: Core assembly and export constituents of each biosynthesis pathway were identified using PFAM domain scanning and BLAST-based homology searches. Based on the genomic locations of core and other neighborhood genes in *M. xanthus*, clusters were generated and their binary distribution was further mapped on to the phylogenetic tree generated via aligning and concatenating thirty housekeeping proteins as shown in this figure. Arrows represent the uninterrupted presence of genes in a cluster. Locus tags highlighted by pale blue boxes correspond to genes such as enzymes involved in monosaccharide synthesis, modification, or incorporation into precursor repeat units of the respective polymer. White circles depict the presence of a homologous gene encoded elsewhere in the chromosome (but not syntenic with the remainder of the EPS/MASC/BPS biosynthesis cluster). Bootstrap values are provided on the tree nodes.

**S2 Figure. (A)** Fruiting body formation phenotypes at different cell densities after growth at 32 ºC for 72 h. **(B)** $EPS^{-}$ (Δ*wzaX*) and $BPS^{-}$ (Δ*wzaB*) cells from exponentially growing cultures were resuspended in buffer to a final concentration of $OD_{600}$ 10. Samples were then spotted on developmental media next to an *E. coli* colony and imaged after 48 h. The first row contains images of the entire swarm; the second row presents a medium-magnification view of the prey invasion step; the third row presents a high-magnification view of the ripples at sites corresponding to the small boxes on the second row.

**S3 Figure. (A)** High-performance anion-exchange chromatography coupled with pulsed amperometric detection for monosaccharide standards vs. monosaccharides isolated from submerged culture supernatants. Strains tested: WT and BPS$^{-}$ (Δ*wzaB*). **(B)** Boxplots of Trypan Blue dye retention to indicate the levels of EPS production in various *pilA* mutant strains relative to WT. The lower and upper boundaries of the boxes correspond to the 25$^{th}$ and 75$^{th}$ percentiles, respectively. The median (line through centre of boxplot) and mean (+) of each dataset are indicated. Lower and upper whiskers represent the 10$^{th}$ and 90$^{th}$ percentiles, respectively; data points above and below the whiskers are drawn as individual points. Asterisks denote datasets displaying statistically significant differences in distributions ($p < 0.05$) shifted higher (*) than WT, as determined via Wilcoxon signed-rank test performed relative to "100" (i.e. WT); stars denote datasets that are not statistically different relative to "0" ($p > 0.05$), as determined via Wilcoxon signed-rank test performed relative to "0" (**S5 Table**). **(C)** Time course of raw surface tension values (via digital drop tensiometry) from representative submerged-culture supernatants. Strains tested: WT, MASC$^{-}$ (Δ*wzaS*), BPS$^{-}$ MASC$^{-}$ (Δ*wzaB* Δ*wzaS*), EPS$^{-}$ MASC$^{-}$ (Δ*wzaX* Δ*wzaS*), EPS$^{-}$ BPS$^{-}$ MASC$^{-}$ (Δ*wzaX* Δ*wzaB* Δ*wzaS*).

**S4 Figure. (A)** Gel chromatography separation of enriched supernatant from a Δ*wzaX* Ω*pilA* culture. **(B)** HSQC spectrum of acidic polysaccharide isolated from Δ*wzaX* Ω*pilA* supernatant. Analysis was performed at 25 ºC, 500 MHz. Resonance peak colours: *black*, C–H; *green*, C–$H_2$.

**S5 Figure. (A)** Dual-labelled ($P_{EPS}$-sfGFP + $P_{BPS}$-mCherry) WT cells (strain EM709) from exponentially growing cultures were spotted on developmental media at a final concentration of

$OD_{600}$ 10.0 and imaged at the indicated time points. Images were scaled as described in Material and Methods. **(B)** Raw, non-normalized data displayed in Panel A

### SUPPLEMENTARY FIGURE 1A

#### EPS cluster

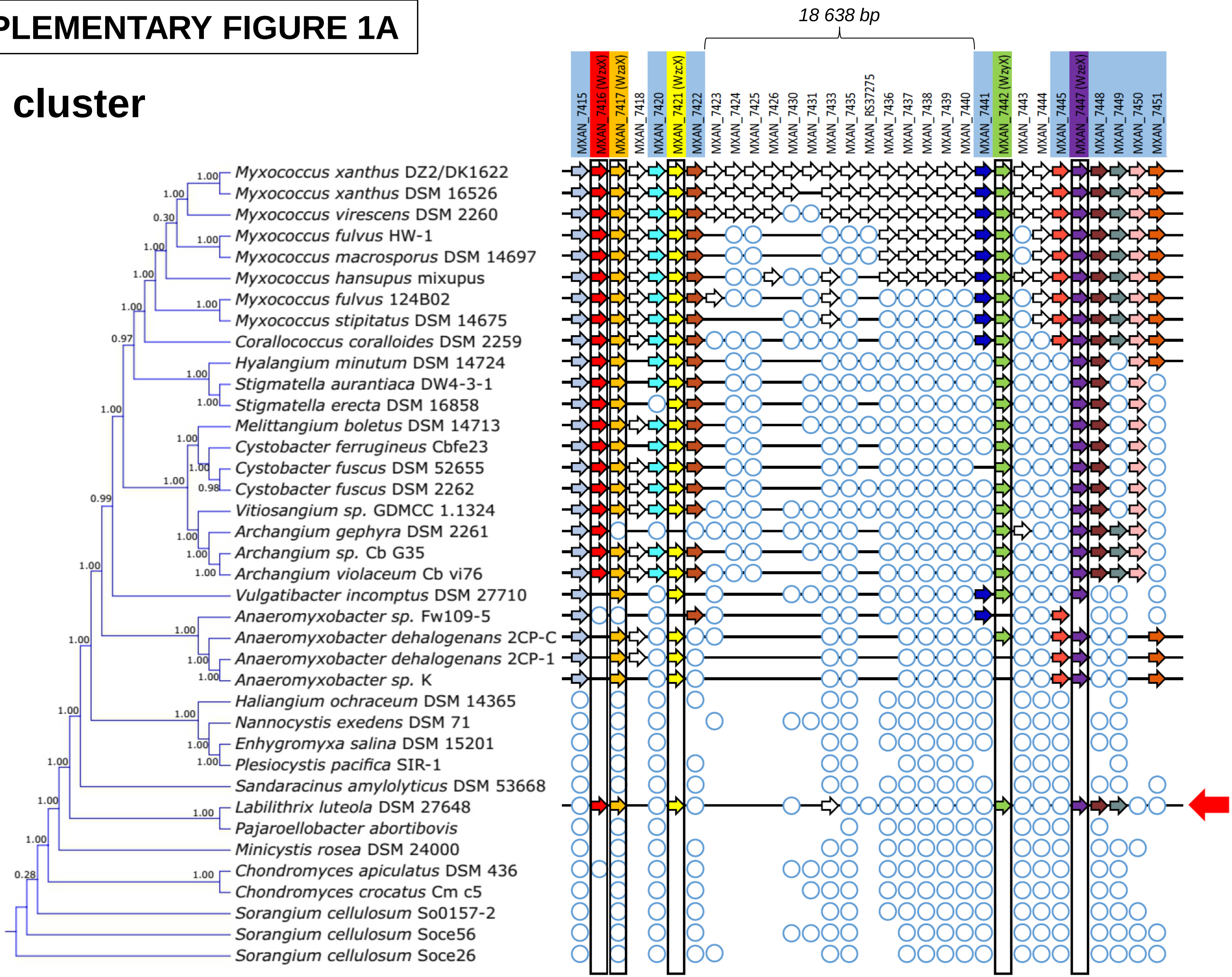

**SUPPLEMENTARY FIGURE 1B**

### MASC cluster

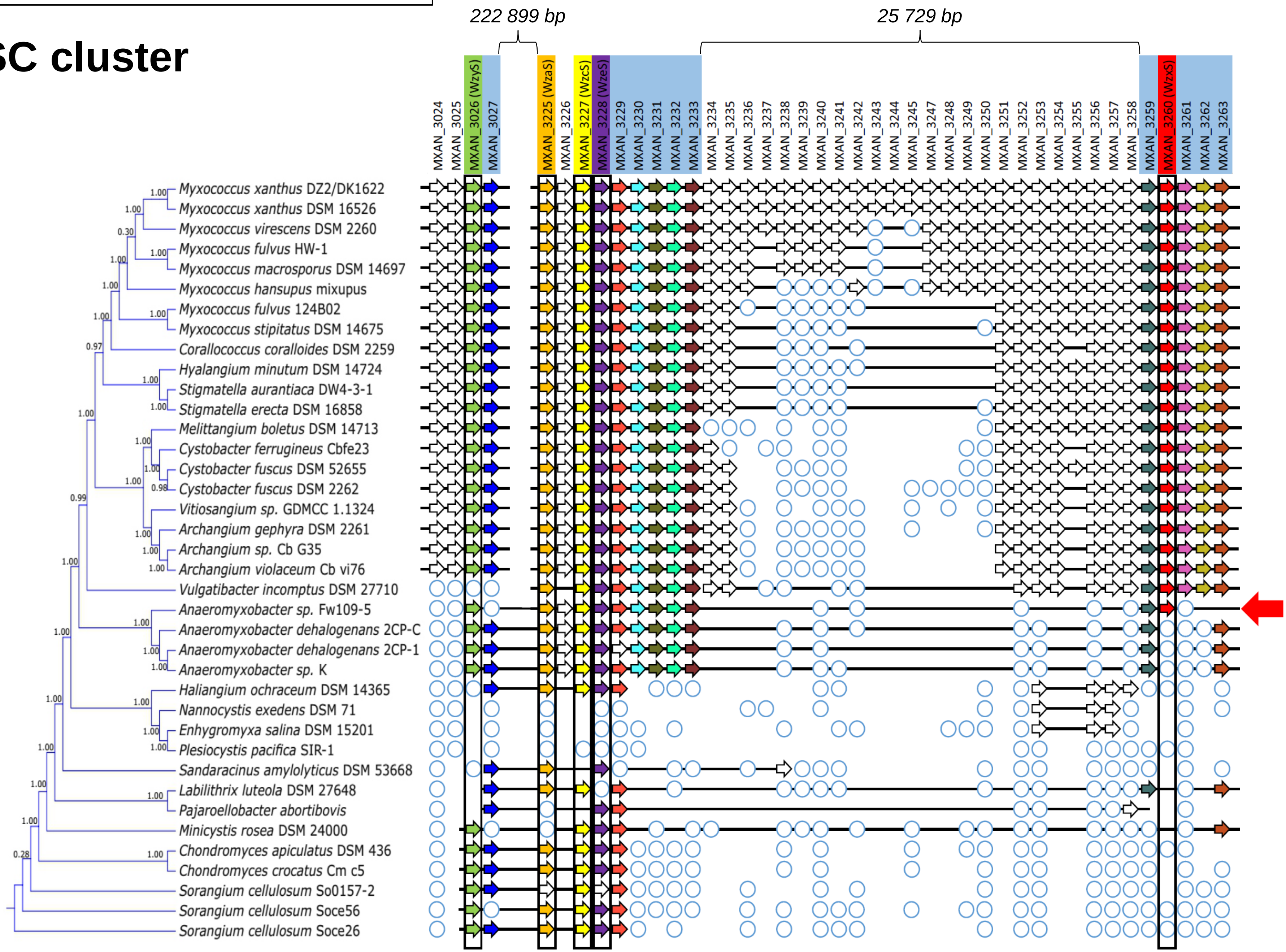

SUPPLEMENTARY FIGURE 1C

### BPS cluster

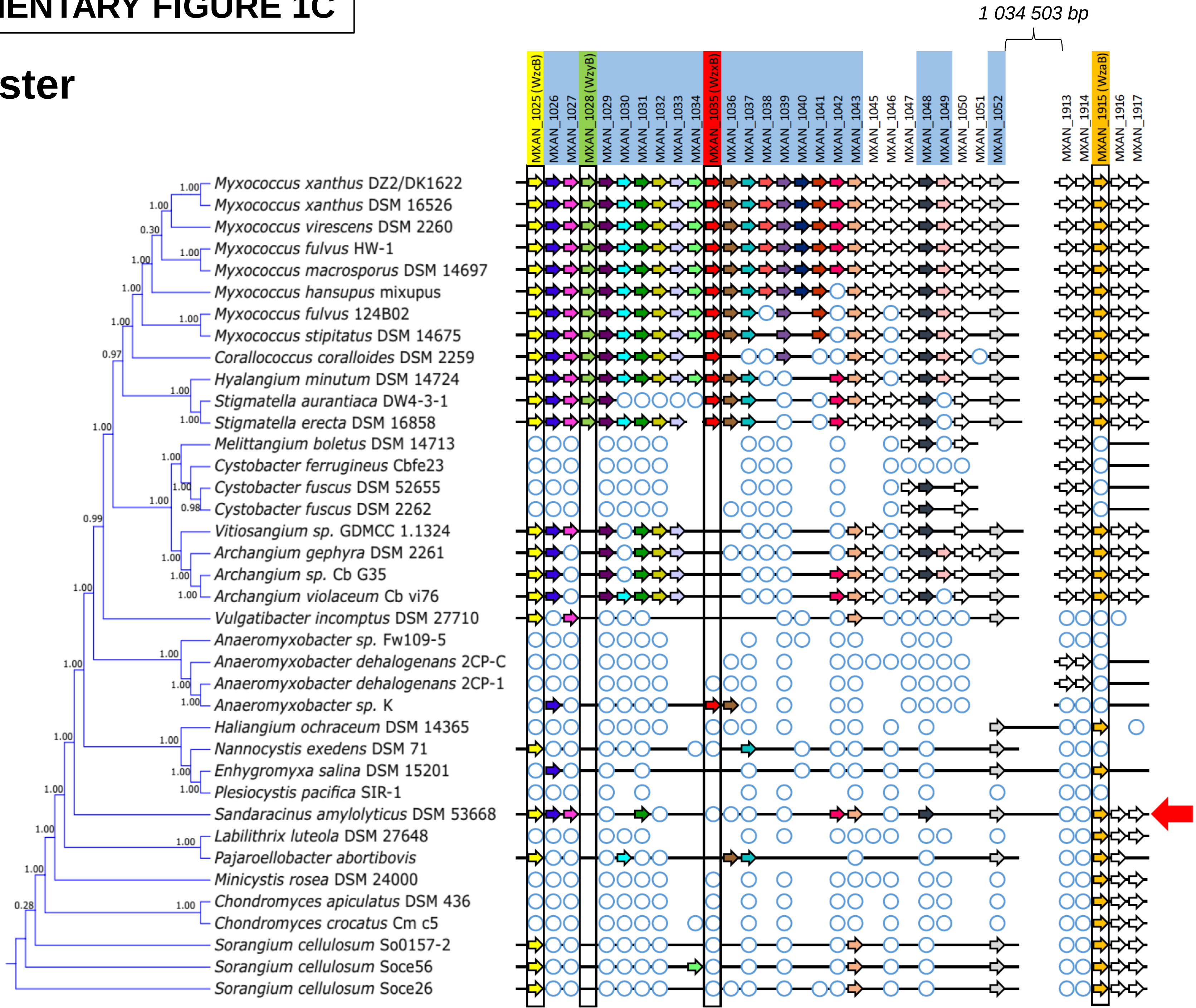

### SUPPLEMENTARY FIGURE 2

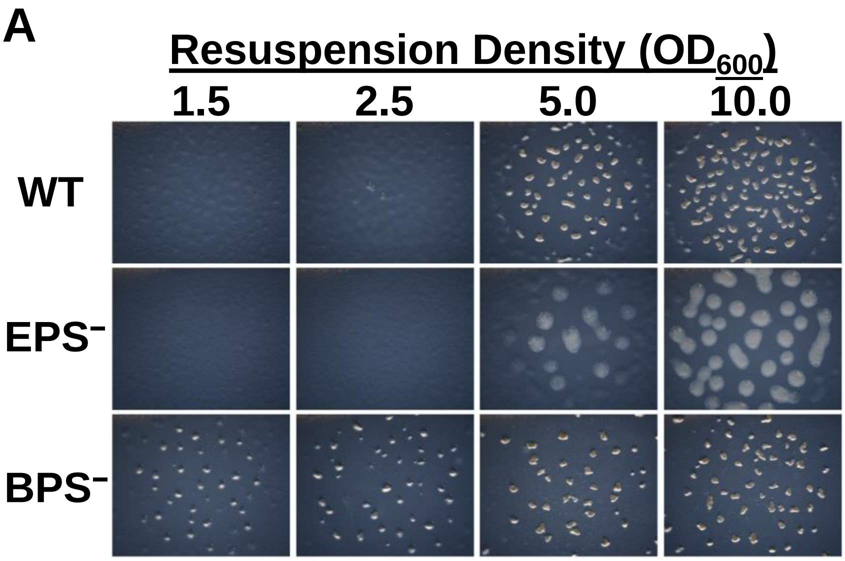


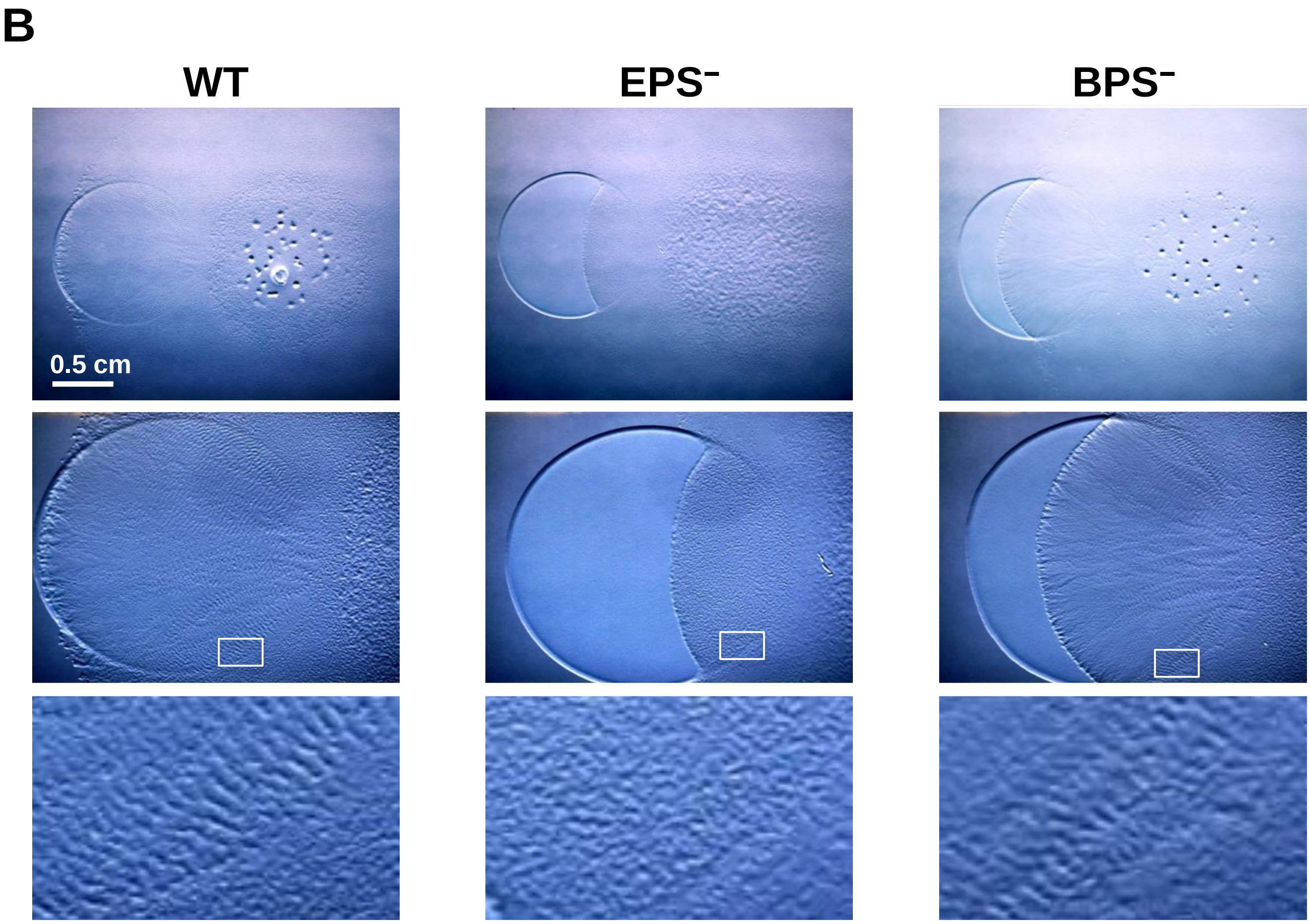

### SUPPLEMENTARY FIGURE 3

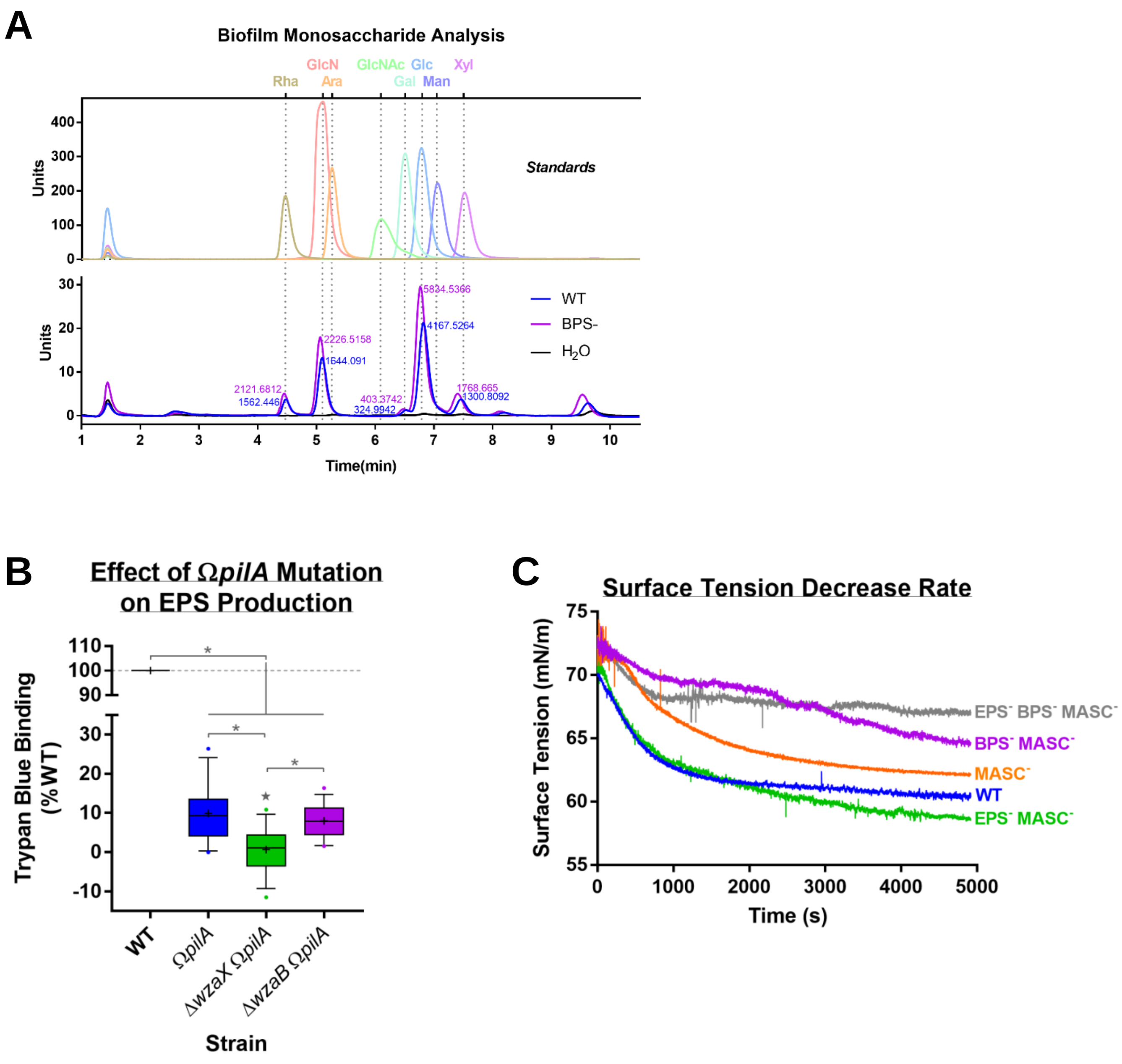

### SUPPLEMENTARY FIGURE 4

**A**

#### *Gel Chromatography*

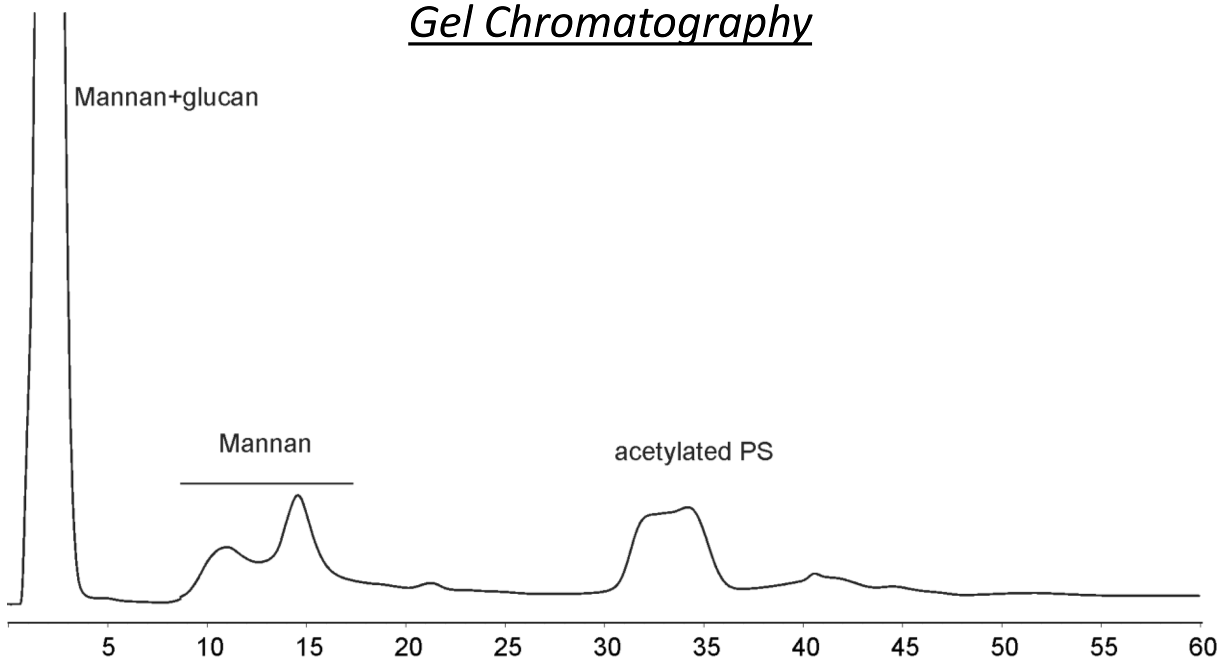


**B**

#### *NMR (HSQC) Spectrum*

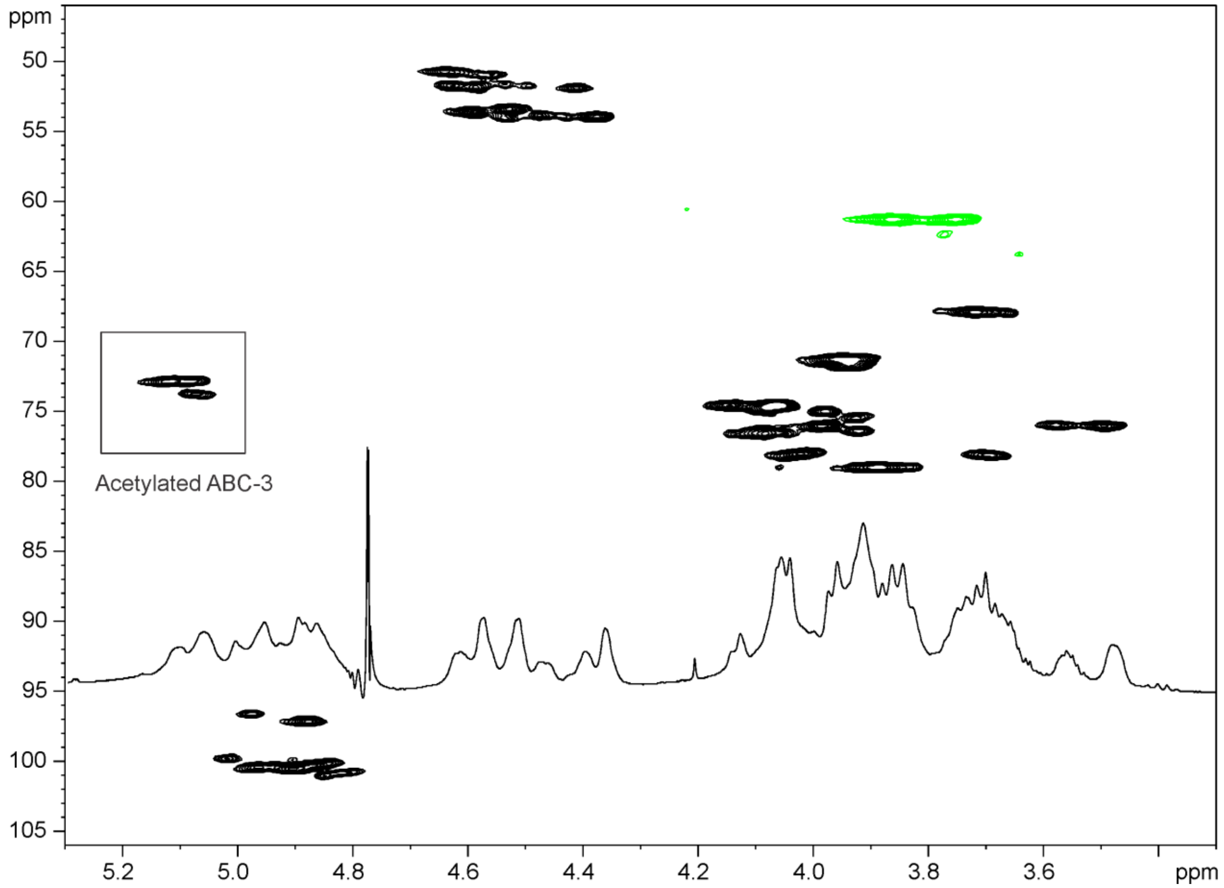

SUPPLEMENTARY FIGURE 5

A

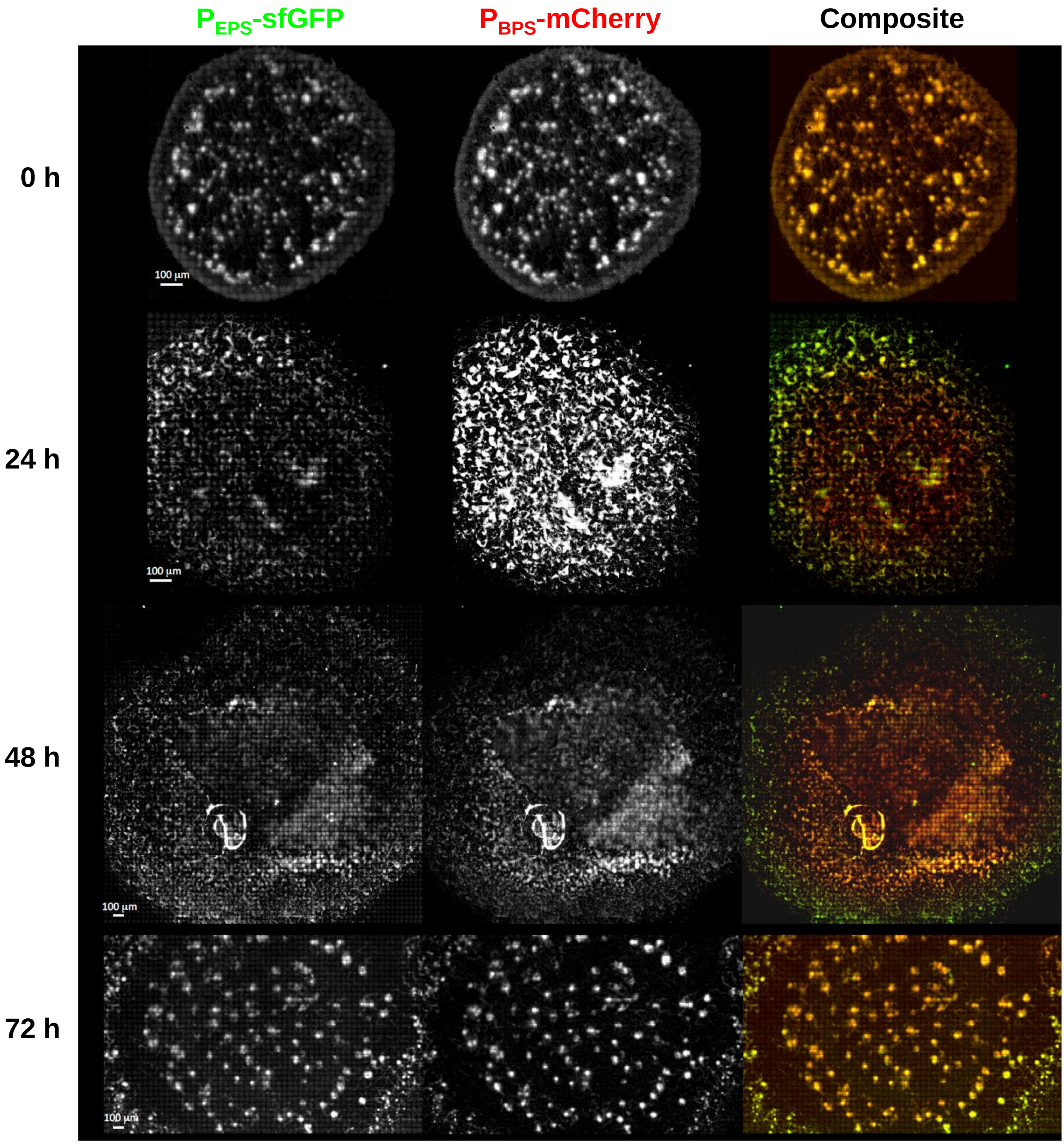

SUPPLEMENTARY FIGURE 5

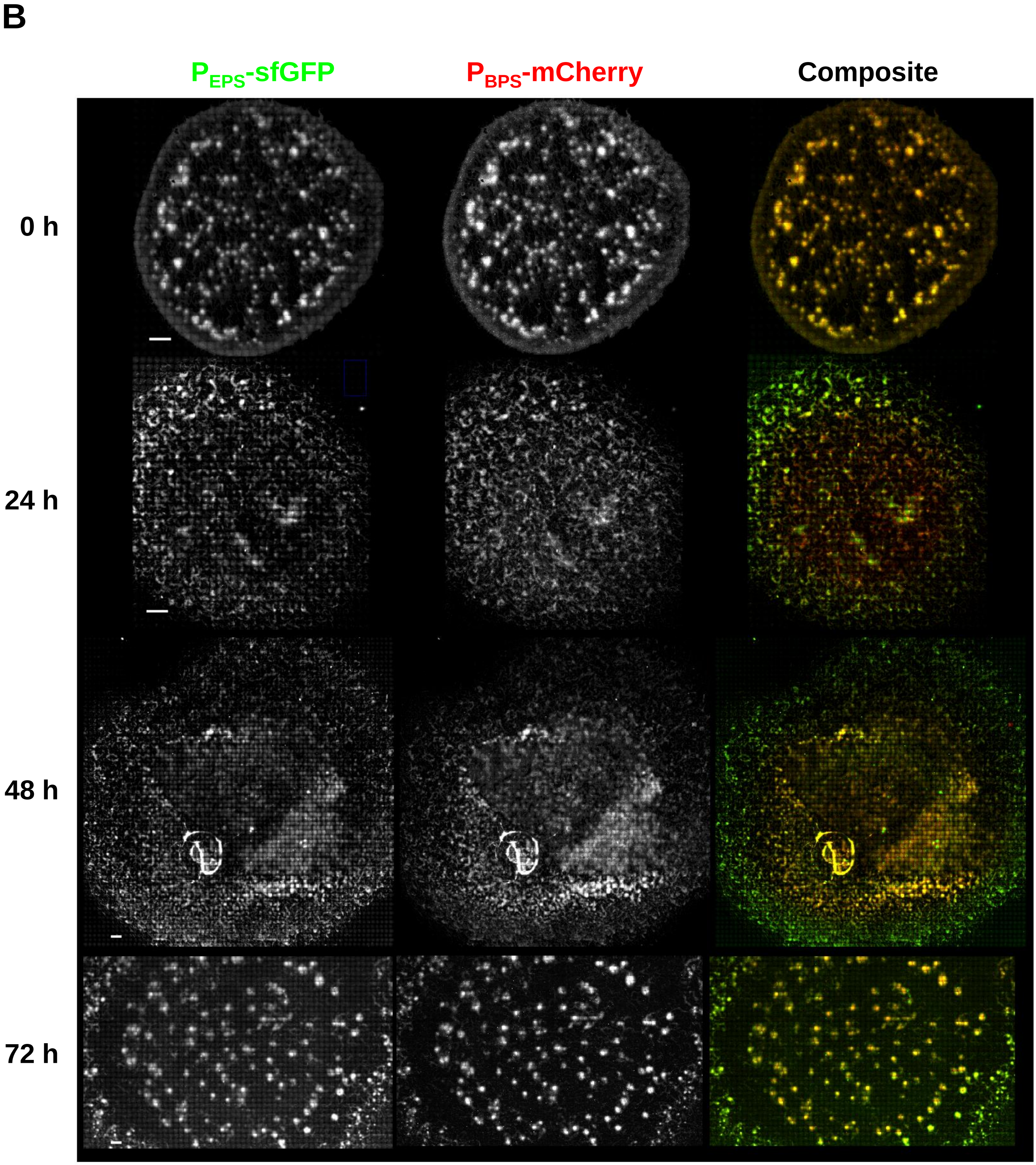
