## Supplementary Table 1 for "Modulation of bacterial multicellularity via spatiotemporal polysaccharide secretion"

**Supplementary Table S1: Bacterial strains and plasmids used in this study**

| Strain Code | Strain | Genotype | Construction | Source or Reference |
|---|---|---|---|---|
| TM108 | *Myxococcus xanthus* DZ2 | Wild type | - | Laboratory collection |
| EM606 | Δ*wzxX* | Δ*mxan_7416* | TM108 + pEM440 (pBJ113-Δ*mxan_7416)* | This work |
| EM616 | Δ*wzyX* | Δ*mxan_7442* | TM108 + pEM470 (pBJ113-Δ*mxan_7442)* | This work |
| EM572 | Δ*wzcX* | Δ*mxan_7421/epsV* | TM108 + pEM441 (pBJ113-Δ*mxan_7421)* | This work |
| EM614 | Δ*wzeX* | Δ*mxan_7447* | TM108 + pEM468 (pBJ113-Δ*mxan_7447)* | This work |
| TM469 | Δ*wzaX* | Δ*mxan_7417/epsY* | TM108 + pTM211 (pBJ114-Δ*mxan_7417)* | Ducret *et al.* 2012 |
| TM484 | Δ*wzaS* | Δ*mxan_3225/exoA/fdgA* | TM108 + pTM210 (pBJ114-Δ*mxan_3225)* | Ducret *et al.* 2012 |
| EM619 | Δ*wzxB* | Δ*mxan_1035* | TM108 + pEM471 (pBJ113-Δ*mxan_1035)* | This work |
| EM618 | Δ*wzyB* | Δ*mxan_1028* | TM108 + pEM472 (pBJ113-Δ*mxan_1028)* | This work |
| EM588 | Δ*wzcB* | Δ*mxan_1025/btkB* | TM108 + pEM462 (pBJ113-Δ*mxan_1025)* | This work |
| EM615 | Δ*wzcB*$_{\mathrm{BYK}}$ | Δ*mxan_1025* $_{\text{bp 1392-2091 (BYK domain)}}$ | TM108 + pEM469 (pBJ113-Δ*mxan_1025* from aa 465-697) | This work |
| TM529 | Δ*wzaB* | Δ*mxan_1915* | TM108 + pTM212 (pBJ114-Δ*mxan_1915)* | Ducret *et al.* 2012 |
| EM596 | Δ*wzcX* Δ*wzaX* | Δ*mxan_7417* Δ*mxan_7421* | TM469 + pEM441 | This work |
| EM592 | Δ*wzaX* Δ*wzaB* | Δ*mxan_7417* Δ*mxan_1915* | TM469 + pTM211 | This work |
| EM591 | Δ*wzcB* Δ*wzaB* | Δ*mxan_1025* Δ*mxan_1915* | TM529 + pEM462 | This work |
| EM651 | Δ*wzaB* Δ*wzaS* | Δ*mxan_1915* Δ*mxan_3225* | TM529 + pTM210 | This work |
| TM488 | Δ*wzaX* Δ*wzaS* | Δ*mxan_7417* Δ*mxan_3225* | TM469 + pTM210 | This work |
| TM530 | Δ*wzaX* Δ*wzaB* Δ*wzaS* | Δ*mxan_7417* Δ*mxan_1915* Δ*mxan_3225* | TM488 + pTM212 (pBJ114-Δ*mxan_1915)* | This work |
| TM293 | Ω*pilA* | Tetracycline resistance cassette | TM108 + *pilA::tet*, Tcr | Laboratory collection |
| TM493 | Δ*wzaX* Ω*pilA* | Δ*mxan_7417* Ω*pilA* | TM469 + Ω*pilA* chromosomal DNA | This work |
| TM540 | Δ*wzaB* Ω*pilA* | Δ*mxan_1915* Ω*pilA* | TM529 + Ω*pilA* chromosomal DNA | This work |
| EM693 | Δ*wzaX* OMss-sfGFP | Δ*mxan_7417 OMss-sfGFP* | TM469 + pSWU19-*promPilA-OMss-sfGFP* | This work |
| EM691 | Δ*wzaB* IMss-mCherry | Δ*mxan_1915 IMss-mCherry* | TM529 + pSWU19-*promPilA-IMss-mCherry* | This work |
| EM709 | $P_{EPS}$-sfGFP + $P_{BPS}$-mCherry | WT $P_{EPS}$*-sfGFP* + $P_{BPS}$*-mCherry* | TM108 + pSWU19-*promWzxX-sfGFP + promWzcB-mCherry* | This work |
| EC393 | *Escherichia coli* MG1655 | $F^{-}$ *lambda*$^{-}$ *ilvG*$^{-}$ *rfb-50 rph-1* | - | Laboratory collection |
