## Supplementary Table 2 for "Modulation of bacterial multicellularity via spatiotemporal polysaccharide secretion"

| **Supplementary Table S2: Gene nomenclature and annotation analyses** | | | | | | | | |
|---|---|---|---|---|---|---|---|---|
| **Gene Name** | **Former name** | **Locus tag** | **Protein length (aa)** | **TMS[1]** | **Fold Recognition Hit (PDB)[2]** | **Name of PDB Homologue** | **Probability (%)** | **E-value** |
| *wzxX* | − | *mxan_7416* | 490 | 14 | 5T77_A | MOP-family lipid II flippase MurJ | 100 | 1.3E-31 |
| *wzyX* | − | *mxan_7442* | 508 | 12 | 6BAR_A | Peptidoglycan glycosyltransferase RodA | 98.66 | 3.3E-6 |
| *wzcX* | *epsV* | *mxan_7421* | 507 | 2 | 4WL1_X | Polysaccharide co-polymerase WzzE | 99.9 | 1.3E-24 |
| *wzeX* | − | *mxan_7447* | 186 | − | 3CIO_D | Tyrosine-protein kinase Etk, C-terminal Wzc domain | 99.41 | 9.1E-14 |
| *wzaX* | *epsY* | *mxan_7417* | 219 | − | 2J58_B | Outer-membrane lipoprotein Wza | 100.0 | 1.3E-32 |
| *wzxS* | − | *mxan_3260* | 506 | 14 | 5T77_A | MOP-family lipid II flippase MurJ | 100 | 2.9E-33 |
| *wzyS* | − | *mxan_3026* | 426 | 12 | 6BAR_A | Peptidoglycan glycosyltransferase RodA | 98.75 | 6.2E-7 |
| *wzcS* | *exoC* | *mxan_3227* | 465 | 2 | 4WL1_X | Polysaccharide co-polymerase WzzE | 99.93 | 7.0E-27 |
| *wzeS* | *exoD, btkA* | *mxan_3228* | 231 | − | 3LA6_D | Nucleotide-binding domain of Tyr-protein kinase Wzc | 99.9 | 9.3E-24 |
| *wzaS* | *fdgA, exoA* | *mxan_3225* | 190 | − | 2J58_B | Outer-membrane lipoprotein Wza | 100.0 | 4.4E-32 |
| *wzxB* | − | *mxan_1035* | 506 | 14 | 5T77_A | MOP-family lipid II flippase MurJ | 100 | 1.8E-33 |
| *wzyB* | − | *mxan_1028* | 486 | 12 | 6BAR_A | Peptidoglycan glycosyltransferase RodA | 98.89 | 2.7E-7 |
| *wzcB* | *btkB* | *mxan_1025* | 710 | 2 | i) aa 2 – 391: 4WL1_X<br>ii) aa 432 – 697: 3CIO_D | i) Polysaccharide co-polymerase WzzE<br>ii) Tyrosine-protein kinase Etk | i) 99.88<br>ii) 99.92 | i) 6.2E-24<br>ii) 6.7E-26 |
| *wzaB* | − | *mxan_1915* | 204 | − | 2J58_B | Outer-membrane lipoprotein Wza | 99.95 | 1.1E-29 |
| [1]Consensus of OCTOPUS and TMHMM analyses. [2]Output from HHpred | | | | | | | | |
